# Understanding Complex Trait Susceptibilities and Ethnical Diversity in a Sample of 4,145 Russians Through Analysis of Clinical and Genetic Data

**DOI:** 10.1101/2023.03.23.534000

**Authors:** Dmitrii Usoltsev, Nikita Kolosov, Oxana Rotar, Alexander Loboda, Maria Boyarinova, Ekaterina Moguchaya, Ekaterina Kolesova, Anastasia Erina, Kristina Tolkunova, Valeriia Rezapova, Olesya Melnik, Olga Freylikhman, Nadezhda Paskar, Asiiat Alieva, Elena Baranova, Elena Bazhenova, Olga Beliaeva, Elena Vasilyeva, Sofia Kibkalo, Rostislav Skitchenko, Alina Babenko, Alexey Sergushichev, Alena Dushina, Ekaterina Lopina, Irina Basyrova, Roman Libis, Dmitrii Duplyakov, Natalya Cherepanova, Kati Donner, Paivi Laiho, Anna Kostareva, Alexandra Konradi, Evgeny Shlyakhto, Aarno Palotie, Mark J. Daly, Mykyta Artomov

## Abstract

The population of Russia consists of more than 150 local ethnicities. The ethnic diversity and geographic origins, which extend from eastern Europe to Asia, make the population uniquely positioned to investigate the shared properties of inherited disease risks between European and Asian ethnicities.

We present the analysis of genetic and phenotypic data from a cohort of 4,145 individuals collected in three metro areas in western Russia. We show the presence of multiple admixed ancestry clusters spanning from primarily European to Asian and high identity-by-descent sharing with the Finnish population. As a result, there was notable enrichment of Finnish-specific variants in Russia. We illustrate the utility of Russian-descent cohorts for discovery of novel population-specific genetic associations, as well as replication of previously identified associations that were thought to be population-specific in other cohorts.

Finally, we provide access to a database of GWAS results for 465 unique phenotypes and allele frequencies.

## Introduction

Linking inherited DNA variation to the disease risks is one of the main goals in modern predictive medicine. Large-scale projects such as the UK Biobank [1], FinnGen [2] and Biobank Japan [3] have made a substantial contribution to the understanding of human biology and the advancement of personalized medicine. The growing ethnic diversity of genetic studies resulted in the discovery of population-specific susceptibility loci that could not be identified with studies limited to a single ancestry [4]. Inclusion of previously underreported populations into biobank initiatives improves fine-mapping accuracy of previously identified GWAS signals and novel risk gene discovery efforts [5].

Genetic studies in the multinational Russian population, which is geographically located at the crossroads of Europe and Asia, could have a potential power to detect historic origins of population-specific variants. Additionally, genetic data from the Russian population could be a powerful source of replication for population-specific associations found in three largest biobanks, UK biobank, FinnGen, and Biobank Japan.

Historically, polygenic trait genetics has been omitted in Russia, resulting in a lack of GWAS studies based on local cohorts. Russian-descent individuals were involved primarily either as a small part of consortia datasets or as a basis for population genetic studies lacking phenotypic information [6-9]. At the same time, there are more than 150 local ethnicities in Russia that would greatly benefit from the local large genetic variation studies. This becomes especially important in light of the lack of transferability of polygenic risk score models between ancestries [10-12].

The aforementioned variety of ethnicities represents the genetic history of populations between Eastern Europeans, Finns, and Asians. For example, a study of whole genome sequencing data previously showed that populations in the northern regions west of the Ural Mountains belong to a Finno-Ugric language group and are genetically close to Finns, while populations in the central regions of western Russia showed similarities with eastern Europeans [13]. Furthermore, close genetic relation between north-western Russians and Finns was also observed in Balto-Slavic speaking populations comparison [14]. In addition, another study showed that Siberian populations separated from other East Asian populations 8,800–11,200 years ago and significantly contributed to the formation of Eastern European populations 4,700-8,000 years ago [15]. Thus, a gene flow from Asia through the Ural Mountains to eastern Europe was hypothesized. Consistently, a notable genetic relationship between Finns and Mongolian tribes was observed [16]. Additional evidence of the great diversity of the Russian population linking European, Asian, and Native American populations was presented in a country-wide study – Genome Russia Project, yet no phenotypic information was collected at that time that impeded biomedical applications [17].

For studies reflecting population history, a small sample set (∼100 individuals) is usually sufficient; however, personalized medicine, biobank assembly and GWAS require a larger sample size and extensive data collection, which is impossible without epidemiological studies. In 2012—2013 in 12 regions of Russia a national study “Epidemiology of cardiovascular diseases in different regions of the Russian Federation” (ESSE-RF) was launched. Within the framework of this study, a stratified multistage random sample of approximately 22,000 residents with blood biobanking and detailed phenotyping was collected [18].

Here, we present the first results of the analysis of clinical and genetic data from three metro areas that participated in the ESSE-RF study: St. Petersburg, Samara, and Orenburg. Our results reflect the genetic structure of the Western Russian population and phenotypic susceptibilities from 4,145 participants across 465 phenotypes.

## Methods

### Data collection

A cohort of 4,800 residents of three areas in Russia – St. Petersburg (N=1,600), Orenburg (N=1,600) and Samara (N=1,600) were recruited in 2012-2013 through an ambulatory visit to local hospitals and polyclinics (**Fig. 1A)**. Detailed phenotypic data and a blood sample were collected. All participants signed an informed consent. The study protocol was approved by local ethics committee at Almazov National Medical Research Center, St. Petersburg.

**Figure 1.**
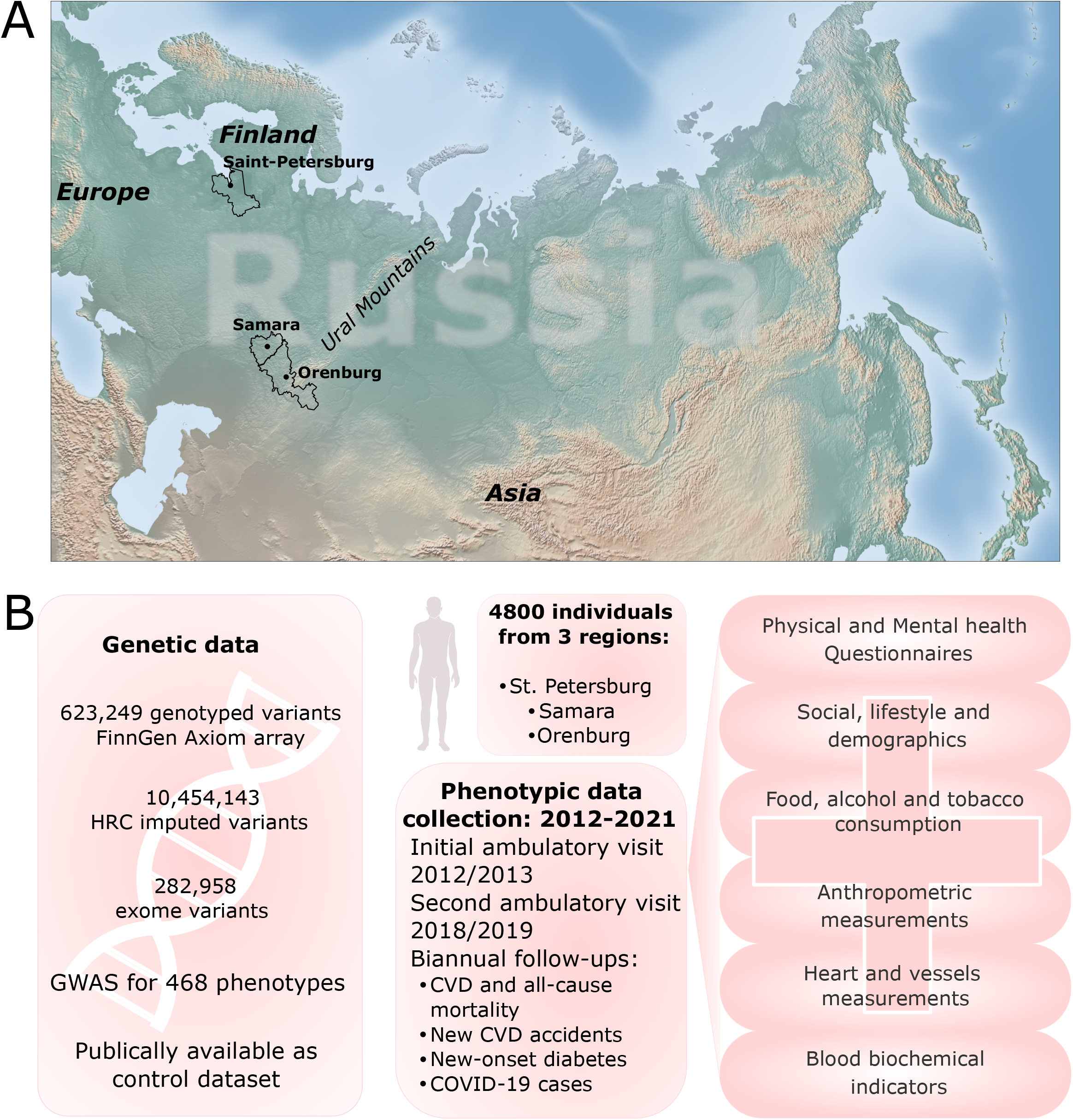
Cohort description and study design. **(A)** Areas in the European part of Russia where sample collection was conducted; **(B)** Ambulatory visit data collection; **(C)** Description of the available genetic, clinical, self-reported and follow-up data.

Each patient was invited for an ambulatory visit for one day to collect phenotypic information (**Fig. 1B)**. Several types of data were collected: responses to the health and medical history questionnaire, dietary behavior, physical activity, social status, depression, anxiety and perceived stress tests, anthropometric parameters, blood pressure and heart rate measurements, blood metabolic panel test, including glucose, creatinine, uric acid, total cholesterol, high-density lipoprotein (HDL), low-density lipoprotein (LDL), triglycerides, insulin and N-terminal prohormone of brain natriuretic peptide (pro-BNP) levels measurements.

For the St. Petersburg cohort, several additional phenotypes were collected: measurements of blood pressure and heart rate in standing position, vessel stiffness measurements and electrocardiogram (ECG), urine albumin and extended blood metabolic panel test with additional test for C-reactive protein (CRP), lipoprotein (a), apolipoproteins A and B, cortisol, leptin, adiponectin, and vitamin D. (**Fig. 1B-C, Sup. Materials, ESSE Data Collection**).

In 2018-2019, 289 out of 1,600 original patients from St. Petersburg were invited for an additional ambulatory visit as a part of different local studies (familial hypercholesterolemia, metabolically healthy obesity, premature vascular aging). These patients were subjected to a detailed follow-up data collection protocol (**Fig. 1C, Sup. Materials, ESSE Follow-up Data Collection**). For all participants, biannual updates on phenotypic and vital status were recorded through direct contact (phone calls) and indirect contacts (mail/e-mail letters, information from local clinical district databases) **Sup. Tab. S1**).

Additional independent cohort of 138 samples were recruited in 2017-2018 for participation as controls in a local study of early childhood starvation effects. They were evaluated using a similar clinical short protocol and no follow-up data collection was performed (**Sup. Materials, Starvation Study Controls Data Collection**).

### Genetic data generation and quality filtering

DNA was extracted from blood samples of 4,723 individuals (4,594 population sample ESSE + 129 Starvation Controls) using the QIAamp DNA Mini Kit (Qiagen) and genotyped using a custom FinnGen ThermoFisher Axiom microarray [2]. All variants and samples with call rate less than 5% were excluded from the study. After initial filtration, additional and missing genotypes were imputed with BEAGLE 4.0 [19] using data from the Haplotype Reference Consortium (HRC) as a reference panel [20]. Directly genotyped and imputed variants with DR2 > 0.8 were included in the downstream analysis [11]. Also, we excluded 252 (224 ESSE and 28 Starvation Controls) samples with mismatch between detected and reported sex.

Relatedness and sample duplications were assessed for individuals using kinship analysis with PLINK2 [21]. 190 duplicates (187 ESSE and 3 Starvation Controls) duplicates were removed and 347 related individuals were labeled for exclusion in further analysis (**Sup. Materials. Kinship analysis. Sup. Fig. S1**). The resulting genetic dataset consisted of 4,281 (4,183 ESSE and 98 Starvation Controls) individuals and 11,077,763 variants. The genetic data was subjected to quality filtration using python3 *‘hail’* (v0.2.85) package [22]. 37,439 variants failing Hardy-Weinberg equilibrium (p < 1×10^−4^) were eliminated from the analyses. Additionally, we removed 371 variants discordant from the HRC imputation panel. The rest of the variants were subjected to comparison of allele frequency against gnomAD Finnish and non-Finnish European populations (**Sup. Materials. Genetic Data Quality Control. Sup. Fig. S2**)

### PCA, population clustering, IBD sharing, and population tree construction

PCA was performed with PLINK2 using 535,727 common autosomal LD-pruned variants (R^2^<0.2). Additionally, we excluded 136 individuals from the ESSE cohort as PCA outliers using *‘adamethods’* (v1.2.1) R package with built-in max Euclidean distances approach [23] (**Sup. Materials. Principal Component Analysis, Sup. Figure S3-S4, Fig. 2A**). Then, we merged the genotyping dataset of ESSE/Starvation Controls with 1000 Genomes WGS data and performed PCA on common 506,617 LD-pruned variants using the python3 *‘hail’* (v0.2.85) package (**Sup. Materials. Comparison of Russian samples with 1000 Genomes, Sup. Fig. S5A, Fig. 2B**). Additionally, we selected ESSE/Starvation Controls and only 1000 Genomes European populations and ran PCA on common 515,649 LD-pruned variants (**Sup. Materials. Comparison of Russian samples with 1000 Genomes, Sup. Figure S5B, Fig. 2C**).

**Figure 2.**
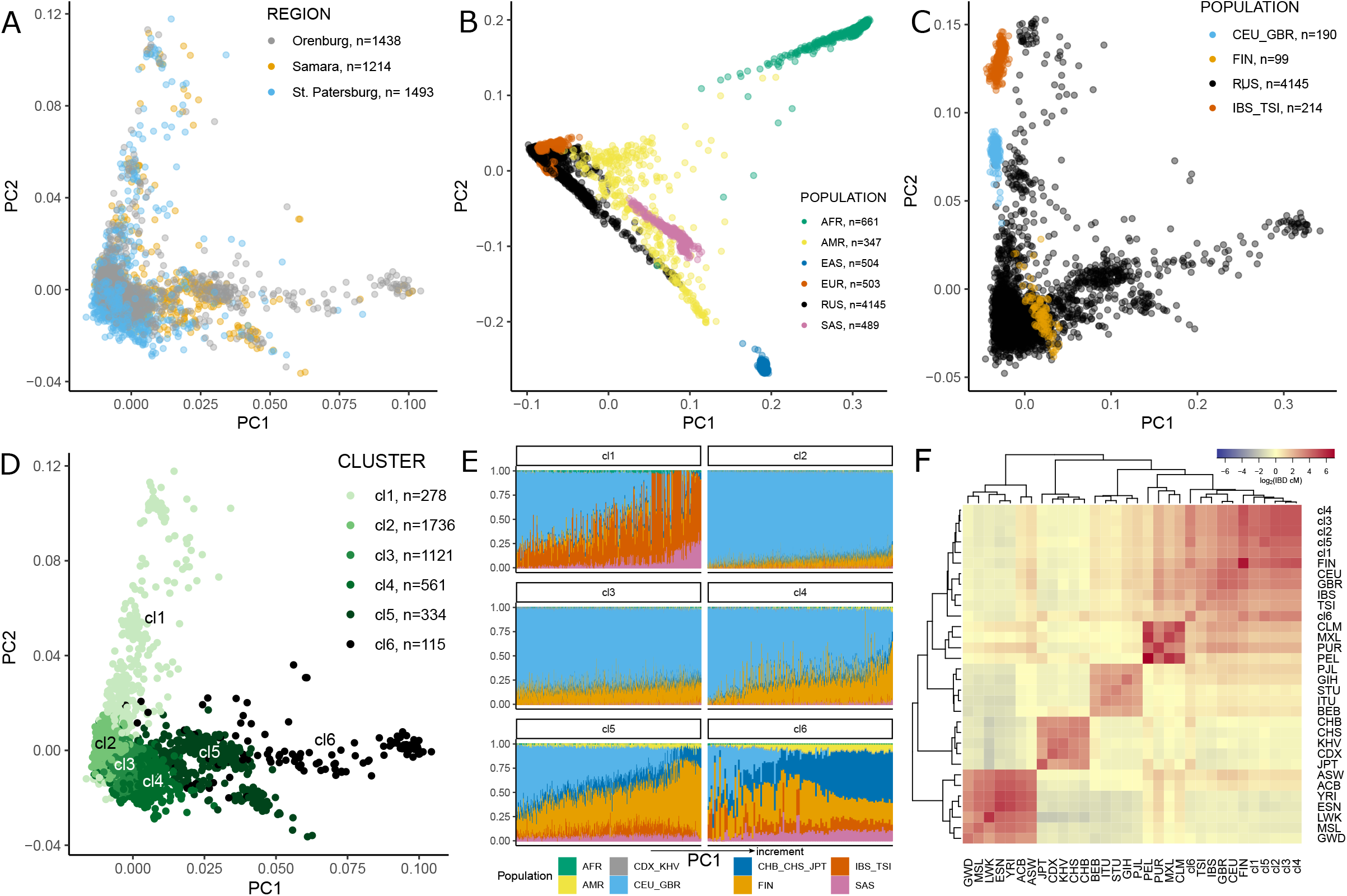
Population structure analysis. Next abbreviations were used: ***AFR*** - Africans (**ACB** - African Caribbean in Barbados, **ASW** - African Ancestry in SW USA, **ESN** - Esan in Nigeria, **GWD** - Gambian in Western Division – Mandinka, **LWK** - Luhya in Webuye, Kenya; **MSL** - Mende in Sierra Leone; **YRI** - Yoruba in Ibadan, Nigeria), ***AMR*** - native Americans (**CLM** - Colombian in Medellín, Colombia, **MXL** - Mexican Ancestry in Los Angeles CA USA, **PEL** - Peruvian in Lima Peru, **PUR** - Puerto Rican in Puerto Rico), ***EAS*** - east Asians (**CHB** - Han Chinese, **CHS** - Han Chinese South, **JPT** - Japanese in Tokyo, **CDX** - Chinese Dai in Xishuangbanna, **KHV** - Kinh in Vietnam), ***RUS*** - Russians, ***SAS*** - south Asians (**BEB** - Bengali in Bangladesh, **GIH** - Gujarati Indians in Houston, Texas, USA; **ITU** - Indian Telugu in the U.K.; **PJL** - Punjabi in Lahore, Pakistan; **STU** - Sri Lankan Tamil in the UK), ***EUR*** - Europeans (**CEU** - central Europeans, **GBR** - British, **FIN** - Finnish in Finland, **IBS** - Iberian populations in Spain, **TSI** - Toscani in Italy). **(A)** Principal component analysis with labelling indicating sample collection region; **(B)** Joint principal component analysis with 1000 Genomes cohort; **(C)** Joint principal component analysis with European subsample of 1000 Genomes cohort**; (D)** Nonoverlapping clustering of the Russian cohort in the PCA space; **(E)** Admixture analysis for each of six populational clusters in the Russian cohort, samples are arranged with respect to their PC1 coordinate**; (F)** Hierarchical clustering of the Russian cohort with 1000 Genomes subpopulations with respect to the sharing of IBD regions.

PCA clusters of Russian samples were identified for 4,145 individuals using *‘SVDFunctions’ (v1*.*2)* R package [24,25]. (**Sup. Materials. Clustering of Russian samples, Sup. Fig. S6, Fig. 2D**). Admixture analysis was performed for combined (ESSE/Starvation Controls - 1000 Genomes) LD-pruned genotype matrix excluding 361 relative individuals (347 - ESSE, 14 - 1000 Genomes). The final dataset for ADMIXTURE (v1.3.0) consisted of 6,649 individuals and 506,617 variants [26] (**Sup. Materials. Admixture analysis, Sup. Fig. S7, Fig. 2E**).

The IBD-sharing statistic was calculated for a combined LD-pruned genotype matrix (ESSE/Starvation Controls - 1000 Genomes) lacking related individuals using BEAGLE 4 (beagle.r1399.jar) (**Sup. Materials. Identity-by-descent (IBD) estimation Fig. 1F, Sup. Fig. S8**). Estimated population size was calculated for resulting IBD regions with length more than 2cM using IBDne (ibdne23Apr20.ae9.jar) tool [27]. Of note, Finnish population size was evaluated only in 1000 Genomes samples, and the small sample size results in larger confidence intervals in the first 10 generations (**Sup. Materials. Estimated population size, Sup. Fig. S9**). Additionally, we calculated F_st_ statistics using VCFtools (v0.1.15) [28] (**Sup. Materials. F**_**st**_ **estimation, Sup. Figure S10-S11**). The population maximum likelihood tree, based on allele counts for each population, was constructed using TreeMix (v.1.12) [29] additional admixture analysis in unsupervised mode was performed using ADMIXTURE (v1.3.0) (**Sup. Materials, Population tree construction**).

### Enrichment of Finnish and Russian variants compared to other European populations

For each variant in the Russian dataset with HWE > 0.0001, we obtained an allele frequency in Non-Finnish Europeans (NFE) and Finnish (FIN) samples from gnomAD. Then, we calculated the log-ratio between allele frequencies in FIN and NFE - the higher it is, the more the corresponding variant is specific to the Finnish population. Only variants passing gnomAD RF filters with call rate >= 0.97 were used in this analysis. We considered 265,624 variants with log-ratio greater than 2 and AF between 0.01 and 0.1 to be Finnish-enriched [2] (**Sup. Fig. S12A**).

For these variants, we calculated the log-ratio between Russian (RUS) AF and gnomAD NFE AF and found that they were also enriched in the Russian population (logistic regression: RUS(1)/NFE(0) ∼ log_2_(AF); p<1×10^−16^, beta=0.15767, se=0.00046); **Sup. Fig. S12B**) [2]. Since the Russian population was divided into 6 clusters, we calculated a log-ratio between AF in each cluster (RUS_cl) and gnomAD NFE AF for all 265,624 Finnish enriched variants. If a variant AF in the cluster was equal to zero, we considered the log-ratio equal to -5.

Additionally, we calculated the log-ratio between gnomAD East Asians AF and NFE AF. If variant AF in the East Asians was equal to zero we considered the log-ratio equal to - 5. Subsequently, all variants with a log-ratio equal to -5 were considered absent in the East Asian population. We defined variants with a log-ratio between RUS AF and gnomAD NFE AF more than 2 as Russian-enriched variants (N=44,963).

### GWAS and other genetic data analyses

Original dataset of 823 phenotypes was subjected to a quality filtration prior to GWAS using R-libraries: *‘dplyr’* (v1.0.0) and *‘tidyr’* (v1.1.1) [30,31]. Only phenotypes with more than 200 individuals were considered for genetic analysis. The final number of phenotypes was 465. Additionally, for 53 continuous phenotypes, we calculated quantile counterparts in which the phenotype was equally divided into 10 quantiles. After that, we filtered the 5th and 95th quantiles of continuous phenotypes to remove outliers. The final GWAS data set included 3,798 unrelated individuals and 7,599,824 variants (MAF>0.01, HWE > 0.0001). As some phenotypes were collected only for a part of the initial cohort, additional quality control was performed (MAF>0.01, HWE > 0.0001) for each phenotype independently. Linear and logistic regression models were used for continuous/categorical and binary phenotypes, respectively. All models were adjusted for sex, age, and PC1-PC4. Some phenotypes were also adjusted for history of medical treatment (**Sup. Materials, GWAS**).

The GWAS results were annotated with VEP [32] and integrated into a PheWeb database [33]. We provide access through an online portal, Biobank Russia: https://biobank.almazovcentre.ru. Furthermore, genetic correlations were calculated between all pairs of phenotypes using LD-score regression, and significant values are shown in the PheWeb database [34,35].

We used POSTGAP [36] and GPrior [37] for gene mapping and prioritization in GWAS to confirm our findings in a case study of smoking status phenotypes. The R *‘ieugwasr’* (v0.1.5) was used to retrieve FinnGen PheWas using the batch ‘finn-b’ [38] for replication. R *‘TwoSampleMR*’ (v0.5.6) was used to clump genome-wide significant Finnish enriched variants [39].

## Results

### Population structure analysis

Initially, we performed PCA to identify ancestral clusters within the dataset (**Methods. PCA, Fig. 2A**). Joint PCA with the 1000 Genomes dataset indicated that the western Russian population is represented by individuals of admixed ancestry, spanning from European to East Asian continental ancestry (**Fig. 2B**). A separate joint PCA with only European subpopulations from 1000 Genomes demonstrated closer relatedness of Russians to Finnish population (**Fig. 2C**). Six clusters within the Russian data set were identified in the PCA space (**Sup. Materials, Clustering of Russian samples, Fig. 2D, Sup. Fig. S5, S6**).

We calculated the contribution of haplotypes from local ancestries represented in 1000 Genomes to each cluster using the ADMIXTURE (**Sup. Materials, Admixture analysis, Fig. 2E, Sup. Fig. S7**). The first cluster included mostly individuals related to southern and eastern Europeans. The second cluster predominantly included a population close to Central Europeans and the British with a small admixture of Finnish and Asian ancestry. In the next four clusters, the proportion of Finnish and Asian haplotypes increases and the proportion of central European haplotypes decreases, which is consistent with the location of these clusters in the principal components space; these clusters co-localized with the Finnish population in PC1-PC2 (**Fig. 2C, Sup. Materials, Comparison of Russian samples with 1000 Genomes)**.

Next, F_st_ and IBD-sharing statistics between Russians and other populations from 1000 Genomes were calculated. We found that according to the F_st_ Russian population in general is close to Central Europeans **(Sup. Fig. S10)**. However, a more precise cluster comparison showed, for example, that cluster 4 is close to Finnish population and cluster 6 is closest to Asian populations which is consistent with PCA and ADMIXTURE analysis **(Sup. Fig. S11, Sup. Materials, Admixture analysis)**.

To verify the relationship between Russians and Finns we calculated IBD-sharing statistics between all Russian clusters and 1000 Genomes populations. We collected all IBD regions with LOD score quality more than 3 for each pair of individuals. Next, we merged the IBD regions if the gap was not greater than 0.6cM and if there was not more than 1 discordant homozygous variant. We summarized the length of resulting IBD regions for each pair of individuals and calculated the median IBD length between pairs of populations (**Sup. Fig. S8**). The resulting heatmap for all populations is shown in **Fig. 2F**. According to the clustering analysis, the Finns are closer to the first five Russian clusters than to the European populations from 1000 Genomes (**Sup. Materials, Identity-by-descent (IBD) estimation)**.

Conclusively, the Russian population sampled in the large metro-areas represents a heterogeneous combination of individuals of admixed ancestries between European and Asian populations.

### Enrichment of Finnish and Russian variants compared to other European populations

The analysis of population structure indicated a complex structure of relatedness between Russians, Finns, and East Asians. The Finnish population is historically unique, however, the genetic similarity of the Russian population illustrated above, suggests that DNA variants enriched in Finnish ancestry might be also found in Russia.

To assess the population-specific properties of DNA variants, we created the distributions of log_2_ allele frequency ratios between the target population and non-Finnish Europeans from gnomAD (NFE) for 265,624 Finnish enriched variants (**Methods. Enrichment of Finnish and Russian variants compared to other European populations; Fig. 3A**). The medians of these log-ratio allele frequency distributions (excluding variants that were not observed) increased from cluster 2 (0.56) to cluster 6 (3.05). In the sixth cluster, it reached an even higher value than in the Finnish population (3.032). As the fraction of East Asian haplotypes increases from cluster 2 to cluster 6 of the Russian population, we investigated the distribution of log-ratios between allele frequencies of Finnish-enriched variants in East Asians and Non-Finnish Europeans (**Fig. 3A, Sup. Tab. S2**). The median value of enrichment was equal to 3.962 which was more than observed in the Finnish population. We found that a significant part (58.35%, N=154,981) of the Finnish-enriched variants have a non-zero frequency in the East Asian population. Furthermore, 45.25%, N=120,204 variants have a log-ratio greater than 2, indicating that these variants are, in fact, more frequent in East Asians than in Finns.

**Figure 3.**
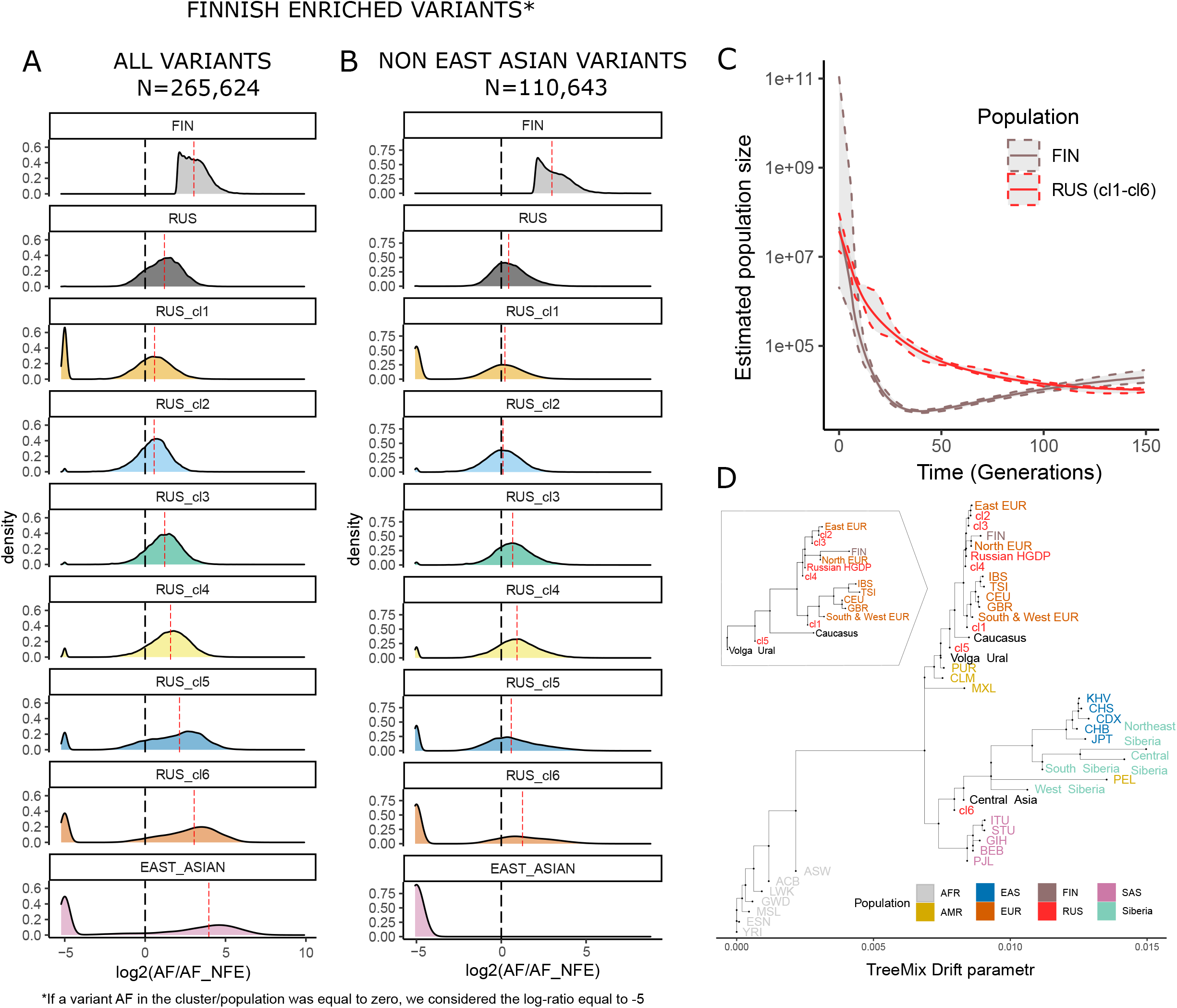
Analysis of prevalence of Finnish-enriched variants in the Russian cohort and TreeMix population structure analysis. **(A)** Distribution of enrichment of the Finnish-enriched variants across clusters in the Russian cohort, Finnish and East Asian cohorts from gnomAD; **(B)** Distribution of enrichment of the Finnish-enriched variants that are not found in East Asian population in gnomAD across clusters in the Russian cohort, Finnish and East Asian cohorts from gnomAD; **(C)** Analysis of estimate population size for Finnish subpopulation from 1000 Genomes cohort and the Russian cohort; **(D)** TreeMix analysis of the relatedness between 1000 Genomes subpopulations, EGDP populations, HGDP Russians and clusters within the Russian cohort.

To investigate the prevalence of Finnish-enriched variants in the Russian sample, we selected only 110,643 variants that were not observed in the East Asian population in gnomAD and found that the highest median enrichments in the Russian cohort were observed in clusters 3-6, which represent the admixed Finno-Asian haplotype structure (**Fig. 3B, Sup. Tab. S2**). Previously reported Finnish-enriched variants associated with clinical phenotypes were generally more prevalent in the Russian population compared to NFE (**Sup. Fig. S13**) [2]. The summary of VEP annotations for Finnish-enriched variants is shown in **Sup. Fig. S14A-B**.

An increase in haplotype frequency usually occurs as a result of the ‘bottleneck’. For example, for the Finnish population several bottlenecks were previously reported [40]. We estimated the population size history for the Russian cohort using Finnish samples from 1000 genomes as a comparison (**Fig. 3C**). No bottleneck was observed in the Russian population, despite the presence of Finnish-enriched variants. We also computed population size based on cluster structure in the Russian population and observed a sign of the bottleneck for clusters with the largest fraction of Asian ancestry (**Sup. Fig. S9**).

We sought to understand the historical origins of the Russian subpopulations and their connection to Finnish and Asian populations, which would potentially explain the presence of population-specific variants. We calculated a maximum likelihood tree using the TreeMix model with reference populations: 1000 Genomes; the populations closest to Russia from the Estonian Genome Diversity Panel (EGDP) [41]; and Russians from the Human Genome Diversity Panel (HGDP) [42]. Interestingly, Finnish samples were the closest to Russians from HGDP and cluster 4 (**Fig. 3D**), while cluster 5 was close to Ural populations (**Sup. Fig. S15**).

Additionally using ADMIXTURE in unsupervised mode with 8 clusters we built a clustering scheme consistent with our tree (**Sup. Fig. S16**). Moreover, we detected a genetic component (shown in yellow in **Sup. Fig. S16**) widely represented in the Siberian population. The fraction of this genetic component decreased with moving from Siberia to the East Europe through the Urals. Also, this component is present in the Finnish population, which may be the result of a known gene flow from Asia to Europe through the Ural Mountains (**Sup. Fig. S16**).

The Russian population also had enrichment of population-specific variants compared to the Europeans, some of which have been described previously [17]. We identified 44,936 Russian-enriched variants as having a log-ratio between RUS and NFE from gnomAD greater than 2. Among Russian-enriched variants 40,743 (90.61%) had higher AF in East Asians than in Russians and only 2,045 (4.55%) were not observed in the East Asian population in gnomAD (**Sup. Fig. 14C-D**).

### GWAS

We performed GWAS for 465 phenotypes and 53 quantile counterparts. Results are available at https://biobank.almazovcentre.ru.

Although 4,600 samples is a relatively modest cohort size for GWAS, we provide examples of replication of findings from other biobanks, as well as newly identified associations specific to the Russian population.

Several associations observed in the UK biobank were replicated - rs7412 and rs4970834 for LDL, rs4697701 and rs4549940 for uric acid levels (**Sup. Fig. S17A-B**).

Interestingly, nominal associations in the UK biobank, such as the association of rs13266066 with the initiation of smoking (UKBB phenocode 20116_0, beta = -0.0043, p=0.00022) [1], later confirmed with the MTAG approach using multiple addiction phenotypes (beta = -0.007, p=1 × 10^−10^, beta was reversed to match models) [43], were significantly associated with the ‘never smoked’ phenotype in the Russian cohort (N never smoked = 2,391; AF never smoked = 0.414, N controls = 1,488; AF controls = 0.475, beta = -0.28, p = 3.74×10-8, **Sup. Fig. S17C**). To reduce the possibility of technical artifacts associated with this observation we looked at only directly genotyped variants in this locus and confirmed the presence of highly-associated rs11781072 (p=1.45×10^−6^). eQTL properties of rs13266066 increased expression of *PTK2* gene in cerebellum (p=8.8×10^−12^). We also performed a gene prioritization analysis which indicated the putative causal role of *PTK2* (**Sup. Fig. S18**).

Several novel genome-wide significant associations for current smoking, abdominal obesity, and increased blood pressure in the second half of pregnancy were identified; however, these would require thorough replication, given a modest discovery cohort size (**Sup. Fig. S17D-F**).

Population structure in the Russian cohort indicated that it could be feasible to use it for replication of Finnish-specific genetic associations. Despite cohort size limitations, we attempted to illustrate this through a systematic approach. First, we selected only potentially replicable (MAF RUS > 0.01) Finnish-enriched variants that were less frequently presented in the East Asian population than in Finns (log-ratio between EAS and NFE < 2). There were 20,050 such variants, which were not clumped together (includes LD-correlates). Among them, we found 142 variants with genome-wide significance associated with 177 traits in FinnGen (total 773 variant-phenotype pairs). Overlaps with the Russian cohort were found for 62 variants in 53 traits (332 variant-phenotype pairs, **Sup. Fig. S19A**).

We combined the phenotypes by similarity into 8 phenotypic groups ‘Diabetes’, ‘Sleep apnea’, ‘Asthma’, ‘Statin’, ‘Alzheimer’, ‘Hypothyroidism’, ‘Arthritis’, ‘Hypertension’, and for each group we independently performed LD-clumping independently (R^2^<0.1) **Sup. Tab. S3**. The resulting data set contained 11 independent variants in 8 phenotypic groups. The only variant rs74800719 passed the Bonferroni-adjusted replication threshold (p=0.05/11 variants/8 traits = 0.00057). This variant was associated with an increased risk of Alzheimer disease in FinnGen (p=1.14×10^−32^, beta= 0.5718) and in the Russian cohort it was associated with an increase in the comorbid phenotype - apolipoprotein B levels (p=4.4×10^−4^, beta=0.15) (**Sup. Fig. S19B**) [44].

## Discussion

The Russian biobank resource presented here is an essential step towards accessibility of precision medicine for the patients with a wide variety of genetic makeups not currently represented in other major genetic studies.

Our cohort illustrates that the genetic structure of the Russian population, sampled in metropolitan areas in the European part of the country, consists of the number of subpopulations with high relatedness to Finnish and East Asian populations. The Finno-Ugric subpopulations in Russia are historically found west of the Uralic mountains, which is in good agreement with previous whole genome studies conducted in this area [9,13]. The gene flow from Asia through Siberia and the Ural Mountains to eastern Europe, together with the previously found relationship between Finns and Asians, suggests that the unique Finnish variants could also be found on the territory of modern Russia. The bottleneck that Finnish population went through resulted in an increase of the allele frequencies of common Finno-Ugric ancestor population. Moreover, our IBD and ADMIXTURE analyses provide potential explanations to high relatedness between Finnish and Mongolian populations reported previously [16].

Replication studies presented here indicate that even with a relatively modest cohort size, previously reported associations from the UK biobank [1] and FinnGen [2] could be directly observed in the Russian cohort. Such a unique genetic structure of the Russian population provides a potential power to discover and replicate associations that often were considered population-specific. Importantly, such replication can be achieved simultaneously with relatively modest cohort size and resources.

This suggests that the susceptibility to polygenic diseases in Russia could potentially be driven by a mixture of variants from multiple ancestral populations. Some of the ancestral populations in this case have not been studied before. Assessing population-dependent contributions of many associated alleles would be critically important for creating informative polygenic risk score models for individual inherited risk evaluation [45].

We continue to monitor participants of this study with the latest data update in the Fall of 2021. Our proof-of-concept study shows that infrastructure, logistics, and research resources are sufficient to create polygenic trait studies in Russia. The major challenge, yet to be resolved, is an outline of how to scale such efforts to the size of other major biobanks.

Finally, we anticipate that the first local resource for polygenic trait genetics studies in Russia, which provides the largest public reference for allele frequencies and genetic associations - Biobank Russia (https://biobank.almazovcentre.ru) – will become a core for further expansion of complex trait genetics research to yet understudied populations.

## Data and code availability

All GWAS, PheWAS, allele frequencies, and aggregated data, along with visualization, are available on the Biobank Russia portal: https://biobank.almazovcentre.ru. The customized pheWeb code is available at https://github.com/ArtomovLab/RusBB_pheweb.

## Author Contributions

M.A., M.J.D., A.P., Al.K., An.K., O.R., E.S. designed the study.

O.R., M.B., E.M., E.K., A.E., K.T., A.D., E.L., I.B., R.L., D.D., O.B., N.P., A.A., E.Bar., E.Baz.,

E.V., S.K., N.C. recruited patients and collected biospecimen.

K.D., P.L., O.M., O.F., An.K. managed the biobank and biospecimen.

D.U., O.R., M.B., E.M., E.K., A.E., K.T. managed and curated the phenotypic data.

D.U., N.K., A.L., V.R., R.S., A.S., M.A. analyzed the data.

D.U., N.K., M.A. designed and created the PheWeb resource.

D.U., O.R., M.A. wrote the manuscript.

M.A. supervised the study.

A.P., M.J.D., An.K., Al.K., E.S., M.A. acquired funding.

All authors reviewed and approved the manuscript.

## Acknowledgements

Genetic data generation and contributions of K.D., P.L., A.P., M.J.D. were supported by the Finnish Academy, Center of Excellence in Disease Genetics

D.U., N.K., O.R., A.L., M.B., E.M., E.K., A.E., K.T., V.R., O.M., O.F., An.K., Al.K., E.S. were supported by the Ministry of Science and Higher Education of the Russian Federation (Agreement # 075-15-2022-301).

The authors thank Dr. Maxim Artyomov (WUSTL), Dr. Manuel Rivas (Stanford University) and Ivan Molotkov (Nationwide Children’s Hospital) for their support of the project and helpful discussions.

