## Supplemental Materials for "Understanding Complex Trait Susceptibilities and Ethnical Diversity in a Sample of 4,145 Russians Through Analysis of Clinical and Genetic Data"

1 - Almazov National Medical Research Centre, Saint-Petersburg, Russia
2 - ITMO University, St. Petersburg, Russia
3 - Broad Institute, Cambridge, MA, USA
4 - Orenburg State Medical University, Orenburg, Russia
5 - Samara State Medical University
6 - Samara Regional Cardiology Dispensary
7 - Institute for Molecular Medicine Finland (FIMM), Helsinki, Finland
8 - Finnish Institute for Health and Welfare (THL), Helsinki, Finland
9 - Analytic and Translational Genetics Unit, Massachusetts General Hospital, Boston, MA USA
10 – Institute for Genomic Medicine, Nationwide Children's Hospital, Columbus, USA
11 – Department of Pediatrics, Ohio State University, Columbus, OH, USA

**Table of contents**


| | |
|---|---|
| **ESSE Data Collection** | 3 |
| **Questionnaire** | 3 |
| **Physical Examination** | 6 |
| **Blood biomarkers** | 7 |
| **Vascular assessment** | 8 |
| **CVD risk scales** | 9 |
| **ESSE Follow-up Data Collection** | 9 |
| **Starvation Study Controls Data Collection** | 10 |
| **Genetic data generation and quality control** | 10 |
| **Kinship analysis** | 10 |
| **Genetic Data Quality Control** | 11 |
| **Principal Component Analysis** | 12 |
| **Comparison of Russian samples with 1000 Genomes** | 13 |
| **Clustering of Russian samples** | 14 |
| **Admixture analysis** | 15 |
| **Identity-by-descent (IBD) estimation** | 16 |
| **Estimated population size** | 17 |
| **Fst estimation** | 18 |
| **Population Tree Construction** | 20 |
| **Enrichment of Finnish variants** | 19 |
| **GWAS** | 22 |

### ESSE Data Collection

A national study "Epidemiology of cardiovascular diseases in different regions of the Russian Federation" (ESSE-RF) was launched in 2012—2013 in 12 regions of Russia, different in climatic, geographic, economic and demographic characteristics. Within the framework of this study, a stratified multistage random sample of 4,800 residents (the adult population - men and women aged 46 ± 12 SD years) was formed in St. Petersburg, Orenburg and Samara. All participants signed informed consent and completed a standard questionnaire based on adapted and validated international methods.

Respondents were invited for a one-day ambulatory visit. First, they filled out a questionnaire, then they underwent anthropometry and donated blood samples. Additionally, in St. Petersburg, electrocardiogram (ECG) was taken and detailed vascular state assessment together with advanced blood tests were performed.

#### Questionnaire

Questionnaire included 10 sections. The first section was about common information of the respondent with contacts, ambulatory visit date, age, sex, birthdate, family status (never married / currently married / divorced / widow), education (primary / incomplete secondary / complete secondary / professional with complete secondary / special secondary / incomplete higher / higher), standing (currently works / never worked / unemployed / retirement pension / disabled) and having children (yes / no and number).

The second section was about dietary behaviors and included questions about salt consumption (always add salt to food / add salt to food before tasting / don't add salt to food), fat usage of cooking food (vegetable oil / margarine / butter / animal fat / no fat at all / mixture of vegetable and animal fats), types of fat used for spreading on bread and/or adding to cereals (none / soft margarine / solid margarine / butter / lard / mixture of vegetable and animal fats), number of lumps and / or teaspoons of sugar (honey, jam, etc.) per day. Also frequencies (I don't use or rarely / 1-2 times per month / 1-2 times per week / daily or almost daily) of red meat, fish or seafood, poultry meat, pickled foods, sausages, pasta or cereals, fresh vegetables and fruits, legumes, sweets and confectionery, milk or kefir or yogurt, sour cream, cottage cheese, cheese were collected. Additionally, the fat content of the used kefir milk, yogurt, sour cream, cottage cheese and cheese was asked. High salt levels were considered as adding salt to food or daily pickled foods consumption. High sugar consumption was considered as ≥ 6 lumps and / or teaspoons of sugar (honey, jam, etc.) per day or daily sweets consumption. High fat consumption was considered as using butter / animal fat for food cooking or using butter / lard for spreading on bread and/or adding to cereals. Insufficient fresh vegetables and fruits consumption was considered as less than daily. Insufficient fish consumption was considered as less than 1-2 times per week.

The third section was about physical activity of respondents and contained 4 questions: 'Which of the following levels of physical activity is most accurate and determines your physical load during work?' (Mostly sit / mostly go / lifting and carrying weights / do hard physical work / don't work). 'How many minutes per day do you walk outside of work, including walking to and from your place of work?' 'How many times a week, in your free time, do you have hard physical activity lasting at least 20-30 minutes such that you have a little shortness of breath or sweat?' How much time, on average, did you spend sitting on a weekday in the last week? Sufficient physical activity was considered as ≥ 150 min walking time per day and hard physical activity ≥ 3 times per week. Low physical activity was considered as < 150 min walking time per day.

The fourth section included questions about smoking status (never smoked / smoked but quit / current smoker), daily smoking (yes / no), age of smoking initiation, smoking quitting and number of cigarettes per day. We calculated the smoking index as a multiplication of the number of cigarettes per day by smoking history (in age) divided by 20. Smoking index ≥ 10 was considered as risky. Additionally, we defined a group currently smokers including those who have not passed more than a year since quitting smoking.

The fifth section included questions about frequency of alcohol consumption (beer, dry wine/champagne, fortified wine, homemade tinctures, spirits), usual amount of drinking at one time and amount of drinking for the last week. Also, this section included an Alcohol Use Disorders Identification Test (AUDIT). Excessive alcohol consumption was defined as ≥ 168g of pure alcohol per week for males and ≥ 84g for females. Additionally, we defined a non-drinking group.

We created combinations between disorder status (obesity / arterial hypertension) and several habitats (high salt / sugar / fat / alcohol / consumption, smoking and low physical activity) for each pair of phenotypes. If the individual did not have a disorder and an unhealthy habit, he was included in the group 0. If the individual had a disorder but hadn't an unhealthy habit, he was included in group 1. If the individual hadn't a disorder but had an unhealthy habit, he was included in group 2. If the individual had both a disorder and an unhealthy habit, he was included in group 3. Additionally, we created a binary factor to compare group 3 with others (0,1,2).

The sixth section included self-assessment scores questions: How do you assess your current state of health? (Scale from 1 to 5). Compared to other people of your age, how do you assess your own health? (Better, same, worse). In this section we also collect the history of menstruation and pregnancy and birth weight: Do you have menstruation? Age of the last menstruation. The reason for the cessation of menstruation (age / artificial menopause with ovaries removing / artificial menopause without ovaries removing). Hormone replacement therapy (yes, no) and hormonal contraception (yes, no). How many pregnancies / births have you had? Have you experienced late fetal loss (after 22 weeks of pregnancy)? Did you give birth before 37 weeks of pregnancy? Have you had an increase in blood pressure in the second half of pregnancy?

The seventh section was devoted to sleep problems and consisted of 7 questions: What was the average duration of your daily sleep over the past month? How often did you find it difficult to fall asleep within 30 minutes after you went to bed? How often did you find it difficult to fall asleep after waking up in the middle of the night? How often have you found it difficult to refrain from falling asleep when the situation calls for it? How often did you have to use medication to fall asleep? Do you snore while asleep? Do you have respiratory arrests during sleep?

The eight section included questions about health history: Do you know your blood pressure / cholesterol levels? Did your doctor tell you that you have high blood pressure / glucose / cholesterol levels? Have you taken any blood pressure / sugar lowering / lipid lowering medications in the past 2 weeks? Do you have headaches regularly? Have you consulted a doctor about headaches? Headaches diagnosis: hypertension, migraine or other. Have you ever experienced sudden short-term weakness or awkwardness when moving in one arm or leg, or arm and leg at the same time? Have you experienced any sudden short-term numbness in one arm, leg or half of your face? Have you ever had a sudden short-term loss of vision in one eye? Have you ever had a sudden dizziness or instability while walking? Has your doctor said that you have a hypertensive crisis? Have you ever received treatment with glucocorticoids or anticonvulsants? Did the doctor tell you that you have / had (osteochondrosis or spondylitis / rheumatoid arthritis / chronic bronchitis / Chronic bronchitis / stroke / myocardial infarction / ischemic heart disease / heart rhythm disorder / heart diseases / diseases of the liver, gallbladder, gastrointestinal tract / stomach ulcer or 12 duodenal ulcer / kidney disease / thyroid diseases / organ transplants / cancer / diabetes mellitus (any type))? Additionally, we asked patients for family history of myocardial infarction, stroke, arterial hypertension, diabetes mellitus (any type) and hip fractures.

The ninth section included 4 questions about the economic conditions of patients: What are the sources of income in your family? What part of your income is usually spent on food? Choose the statement that most accurately describes the financial capabilities of your family (We don't even have enough for the basic necessities / We can buy everything we need, but we can't buy expensive durable / From time to time we may purchase expensive goods durable / We can buy durable goods, but we can't buy things like an apartment, a house, or an expensive car / We are able to buy things like a house, an apartment or an expensive car). How do you rate your family's well-being compared to others? (Very wealthy / relatively wealthy / average / relatively poor / very poor).

The last section includes a questionnaire assessing perceived stress (PSS) [1]. The Hospital Anxiety and Depression Scale (HADS) was used to assess depression and anxiety [2].

The Rose questionnaire was used as a screening test for the diagnosis of Angina pectoris [3]. The standard questionnaire consists of 9 questions that identify predictors of angina. Questions 10, 11 and 12 have been added by the Federal State Budgetary Institution National Medical Research Center for Therapy and Preventive Medicine (department of multifactorial prophylaxis) [4]. Also, intermittent claudication and risk of heart attack were assessed.

#### Physical Examination

All participants underwent anthropometry. The measurement of body weight was carried out on the scales of the brand VEM-150- Mass-K (Russia), height — using the height meter RM-1 Diacoms (Russia), waist and hip circumference — using a standard flexible centimeter tape. Abdominal obesity was determined according to the JIS 2009 metabolic syndrome criteria 2009 [5] as: waist circumference (WC) ≥94 cm for males and ≥80 cm for females. Additional less strict criterion was ≥ 102 cm for males and ≥88 cm for females. Obesity based on waist to hip circumference ratio was defined as ≥ 0.9 for males and ≥ 0.85 for females. Body mass index (BMI) was calculated using the Quetelet formula as the ratio of body weight in kilograms to height in meters squared. All respondents were classified as obese (BMI ≥ 30 kg/m2), overweight (25 kg/m2 ≤ BMI <30 kg/m2), normal weight (18.5 kg/m2 ≤ BMI < 25 kg/m2) and underweight (BMI < 18.5 kg/m2). Additionally, obese group (BMI ≥ 30 kg/m2) was separated into 3 subgroups according to the degree of obesity: 1 degree (30 kg/m2 ≤ BMI <35 kg/m2), 2 degree (35 kg/m2 ≤ BMI <40 kg/m2), 3 degree (≥40 kg/m2). Finally, we defined groups of individuals who had at least 1 type of obesity (abdominal, waist to hip circumference ratio or BMI).

Blood pressure (BP) and heart rate (HR) were measured by the OMRON tonometer (Japan) after resting for 5 minutes in a sitting position twice on the right (N=4,339) or left (N=457) hand with intervals of 2 minutes. The average BP and HR of the two measurements was calculated. Antihypertensive treatment in the past 2 weeks was clarified through a questionnaire. Two cutt-off levels were applied to BP: ≥ 140/90 mmHg according to hypertension guideline [6] and ≥ 135/80 mmHg according to metabolic syndrome consensus [5]. All responders classified as having arterial hypertension (AH) (BP ≥ 140/90 mmHg or antihypertensive treatment), high-normal BP (135/85 mmHg ≤ BP <140/90 mmHg, no antihypertensive treatment) normal BP (120/80 mmHg ≤ BP <135/85 mmHg, no antihypertensive treatment) and optimal BP (BP < 120/80 mmHg, no antihypertensive treatment). High-normal BP and normal BP was considered as prehypertension. Efficient antihypertensive treatment was defined as having BP < 140/90 mmHg and more strict criteria as having BP < 130/80 mmHg [6].

We calculated the Estimated Pulse Wave velocity as it was defined previously [7]. Mean blood pressure was calculated as DBP + 0.4*(SBP - DBP). Responders from St. Petersburg additionally had orthostatic BP and heart rate measurement after 3 minutes in standing position.

Electrocardiogram (ECG) registration was carried out using a computerized PADSY complex (Medset Medizintechnik GmbH, Germany), interpretation was carried out according to a standard protocol.

#### Blood biomarkers

*3 areas (St. Petersburg, Samara, Orenburg)*

The blood glucose (mmol/L), creatinine (µmol/L), uric acid (µmol/L.), and lipids (mmol/L) measurement (total cholesterol – TC, low-density lipoproteins – LDL, high-density lipoproteins – HDL, triglycerides – TG) were performed in fasting state (Abbott Architect 8000, USA; reagents Abbott Diagnostic). We defined several groups of respondents according to glucose levels: 1st group (glucose < 5.6 mmol/L), 2nd group (5.6 mmol/L ≤ glucose < 7 mmol/L), 3rd group (7 mmol/L ≤ glucose < 11 mmol/L) and 4th group (glucose ≥ 11 mmol/L). Glucose lowering and lipid lowering therapy in the past 2 weeks were clarified through a questionnaire. Dyslipidemia was considered for individuals with increased cholesterol > 4.9 mmol/L or LDL > 3 mmol/L or TG > 1.7 mmol/L or reduced HDL (in male <1.0 and in female <1.2 mmol/L) or presence of lipid-lowering therapy [8]. Additionally, the atherogenic index of plasma (AIP) was calculated as logarithmic ratio between triglycerides and HDL: $\log_2$(TG/HDL). The glomerular filtration rate (GFR) was calculated using the CKD-EPI formula [9]. Diabetes was diagnosed in case of anamnesis (information from patient).

Insulin (pmol/L) and N-terminal pro-hormone of brain natriuretic peptide (pro-BNP, pg/ml) levels were performed in fasting state (Cobas Integra 400 plus, Switzerland; Roche-diagnostics reagents). Two cutt-off levels were applied to pro-BNP (pro-BNP >125 pg/ml and pro-BNP > 300 pg/mL) [10]. Insulin resistance index was calculated using the formula: fasting blood glucose × insulin × 0.138 (coefficient used to convert pmol/L to µU/ml) / 22.5 (HOMA-IR (Homeostasis Model Assessment of Insulin Resistance)). HOMA-IR was defined in the case of Insulin resistance index ≥ 2.6. The triglyceride glucose (TyG) index was calculated as a natural logarithm: ln(TG[mmol/L] × 87.5 × glucose[mmol/L] × 18/2). Modifications of the TyG index were calculated as TyG_BMI = TyG × BMI, TyG_WC = TyG × Waist circumference and TyG_WC_HEI = TyG × Waist circumference/Height [11]. Low insulin levels were considered as < 17.8 pmol/L, high as > 173 pmol/L. The FINDRISC scale was used to assess risks of diabetes [12].

Additionally, the metabolic syndrome was determined according to the following criteria (presence of three or more components: SBP ≥130 or DBP ≥85 mm Hg or antihypertensive therapy; triglycerides ≥ 1.70 mmol/L; HDL <1.04 (males)/1.30 (females) mmol/L or lipid-lowering therapy; glucose ≥5.6 mmol/L or hypoglycemic therapy; WC >102 (males)/88 (females)) in combination with the absence of cardiovascular diseases (CVD) and diabetes mellitus (DM) at the time of inclusion in the study, according to the anamnesis [13].

*St. Petersburg only (additional blood biomarkers)*

The leptin and adiponectin levels were determined by enzyme-like immunoassay (DRG, Germany). C-reactive protein (CRP,mg/L), thyroid-stimulating hormone (TSH,mIU/L) and cortisol (μmol/L) were measured in fasting blood samples (Cobas Integra 400 plus, Switzerland; Roche-diagnostics reagents). Two cut-off levels of CRP were applied: CRP > 3 mg/L and CRP > 1 mg/L [14]. TSH > 4 mIU/L was defined as a high TSH level [15,16]. Low TSH was considered as < 0.4 mIU/L [15]. Low cortisol was defined as < 171 μmol/L.

Extended lipids study (lipoprotein (a), apolipoprotein A, apolipoprotein B) and urine microalbuminuria (albumin excretion in urine portion) was performed using (Abbott Architect 8000, USA; reagents Abbott Diagnostic). High lipoprotein (a) was considered as > 0.3 g/L [17]. Low apolipoprotein A was defined as < 1.04 g/L for male and < 1.08 g/L for female. High apolipoprotein A was defined as > 2.02 g/L for male and > 2.25 g/L for female. Low apolipoprotein B was defined as < 0.66 g/L for male and < 0.6 g/L for female. High apolipoprotein B was defined as > 1.33 g/L for male and > 1.17 g/L for female. High urine albumin excretion was defined as > 30 mg/dL [6].

#### Vascular assessment

Vascular assessment was performed only for St. Petersburg residents.

Ultrasound examination of the common carotid arteries was performed using a portable diagnostic system My Sono U6 (Samsung, Korea). The standard protocol included measurements bilaterally at a distance of 1 cm from the bifurcation of the common carotid artery along its posterior wall in three positions (anterior, middle, and posterior longitudinal). The thickness of the intima-media complex (IMT) was defined as the distance between the first and second echogenic line of the located vessel. Subsequently, the mean IMT on both sides was calculated as the arithmetic mean of three measurements. In addition, the presence or absence of atherosclerotic plaques was assessed. Values greater than 0.9 mm and less than 1.3 mm were taken as an increased IMT. Local thickening ≥ 1.3 mm was regarded as atherosclerotic plaques [18].

The carotid-femoral pulse wave velocity (cfPWV) was assessed using a SphygmoCor device (AtCor, Australia). Carotid-femoral distance was measured using the formula recommended by the 2012 Expert Consensus on Vascular Stiffness: (distance from common carotid artery to common femoral artery in cm) × 0.8. Using a special sensor, the applanation method recorded pulse waves for 10 seconds, first in the projection of the common carotid artery on the left, then in the common femoral artery on the left, also for at least 10 seconds. An indicator of less than 10 m/s for PWV was taken as the normal value [19].

Cardio-ankle vascular index (CAVI) was measured automatically on the right and left using the VaSera VS-1500 device (Fukuda Denshi, Japan). CAVI was calculated between the heart valve and the ankle artery using the FCG signal (II tone) and plethysmographs obtained by applying cuffs to the upper arm and lower leg. The value of the CAVI > 9.0 was considered as elevated [20]. Measurement of ankle-brachial index (ABI) was also performed automatically on the right and left, calculated as the ratio of SBP on the leg (at the ankle) to SBP on the brachial bilaterally artery. ABI less than 0,9 at least one side was considered decreased [6].

#### CVD risk scales

Several cardiovascular risk scores were calculated using well-known scales. Framingham risk score 2008 calculated using mean parameters values obtained from the ESSE cohort and from the original study [22] and using Framingham risk score 2008 tables. ASCVD risk estimator 2013 calculated with mean parameters values obtained from ESSE cohort and from the original study [23]. SCORE systemes 2003, 2016 and 2019 were calculated [24,25,26]. The SCORE system predicts only fatal cardiovascular disease. To convert fatal risk to total (non-fatal + fatal) we multiplied SCORE-High ASCVD risk results to 3 for men and to 4 for women [8]. The recalibration for the Russian population scale of SCORE 2016 - SCORE-MoSP was also calculated [27].

### ESSE Follow-up Data Collection

Additionally, we collected data about patients' vital status, cardiovascular events and new disease onset each two years (**Sup. Tab. S1).** Biennially from 2013 to 2021, participants and their relatives were contacted by phone and e-mail to collect information regarding fatal events (all-cause mortality) and non-fatal events (myocardial infarction, unstable angina, stroke). Patients and their relatives were invited to the clinic where information was verified in medical documentation and death certifications.

289 patients from St. Petersburg were invited for detailed follow-up data collection visits in 2018-2019 as part of different local studies (familial hypercholesterolemia, metabolic healthy obesity, early vascular aging). We collected the same phenotypic information as for the first visit using the analogous protocols and methods (**Sup. Materials, ESSE Data Collection**). For all continuous phenotypes we calculated delta as the difference between the first and second visit measurements.

### Starvation Study Controls Data Collection

138 individuals were recruited in 2017-2018 in St. Petersburg as controls for local study of early childhood starvation effects. Respondents were invited for a one-day ambulatory visit.

Initially they filled out a short version of the ESSE questionnaire with questions about family status, education, smoking status, antihypertensive / glucose lowering / lipids lowering therapy and concomitant diseases. Anthropometry, blood pressure and heart rate in sitting and standing positions were collected using the same methods as for ESSE respondents (**Sup. Materials, ESSE Data Collection**). Blood samples were collected and biobanked. Fasting glucose, creatinine, and lipids were performed (**Sup. Materials, ESSE Data Collection**).

### Genetic data generation and quality control

4,723 individuals (4,594 ESSE + 129 Starvation Controls) were genotyped using a custom FinnGen Affymetrix Axiom array [28]. After the genotype imputation with BEAGLE 4.0 [29] using Haplotype Reference Consortium (HRC) data as a reference panel [30] the resulting dataset contained 623,249 genotyped variants and 10,454,514 imputed variants. We exclude 252 (224 ESSE and 28 Starvation Controls) samples with sex mismatch.

#### Kinship analysis

We used PLINK2 [31] to identify relationships between individuals in the Russian population. We defined the 1st relationship degree as kinship value between 0.177 and 0.354 and second degree as value between 0.088 and 0.177. Thus, we had 400 pairs of the first degree relationship and 66 pairs of the second degree. We used *'igraph'* (v 1.2.11) R [32,33] package to visualize all relatives (**Sup. Fig. S1**). Then using PLINK2 kinship estimator we marked 347 individuals among relatives in such a way that when they were excluded, only unrelated ties remained. Also, we identified 190 individuals with kinship more than 0.354 which correspond to twins. We defined these samples as duplicates and removed them from the study.

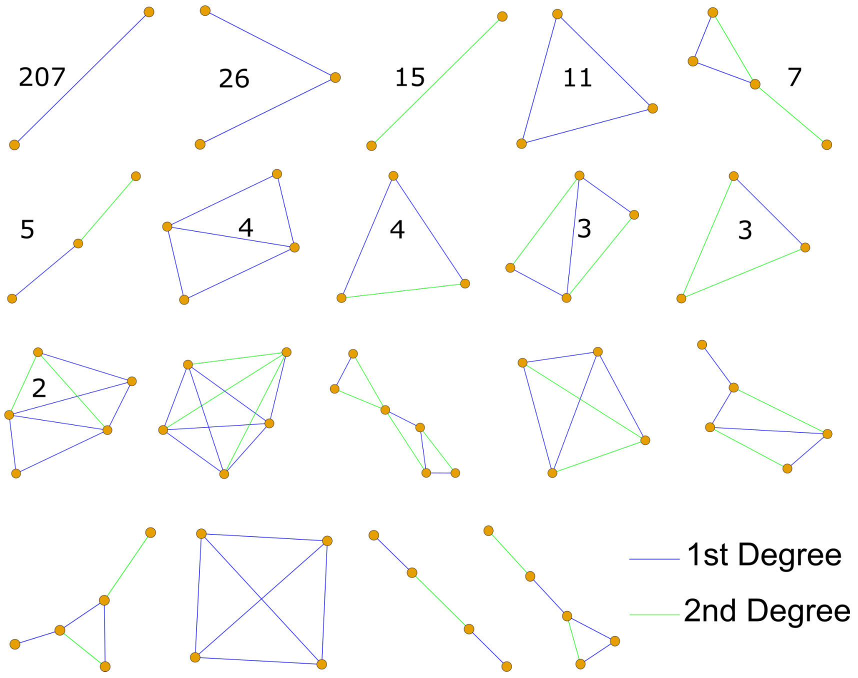


**Supplementary figure S1.** All sets of relative groups in the combined ESSE-Starvation controls dataset. Only among ESSE cohort relatives were detected.

#### Genetic Data Quality Control

We used python3.6 *'hail'* (v0.2.85) [34] package to assess the variant quality control. Initially we removed 37,439 variants failing Hardy-Weinberg equilibrium ($p < 1x10^{-4}$) and 371 discordant variants with the HRC imputation panel. AF of discordant variants differed from the HRC panel more than 0.15.

The distribution of MAFs and allele count for rare variants (MAF<0.01) for directly genotyped and all variants are shown at **Sup. Fig. S2A-B,** correspondingly. Majority of rare genotyped variants had 0 allele count, while the full set of rare variants predominantly had 1-20 allele counts (**Sup. Fig. S2A-B)**.

Next, we compare MAF of our Russian variants with MAFs from non-Finish and Finnish gnomAD samples. We used gnomAD hail matrix and filtered all variants with call rate less than 0.97. Also, we filtered out all variants that did not pass gnomAD Random Forest filters. Therefore only 537,363 genotyped variants and 10,049,642 imputed variants were included into comparison. Comparison with non-Finish Europeans for genotyped and all variants is shown at **Sup. Fig. S2C-D**, correspondingly. Comparison with Finish for genotyped and all variants is shown at **Sup. Fig. S2E-F**, correspondingly.

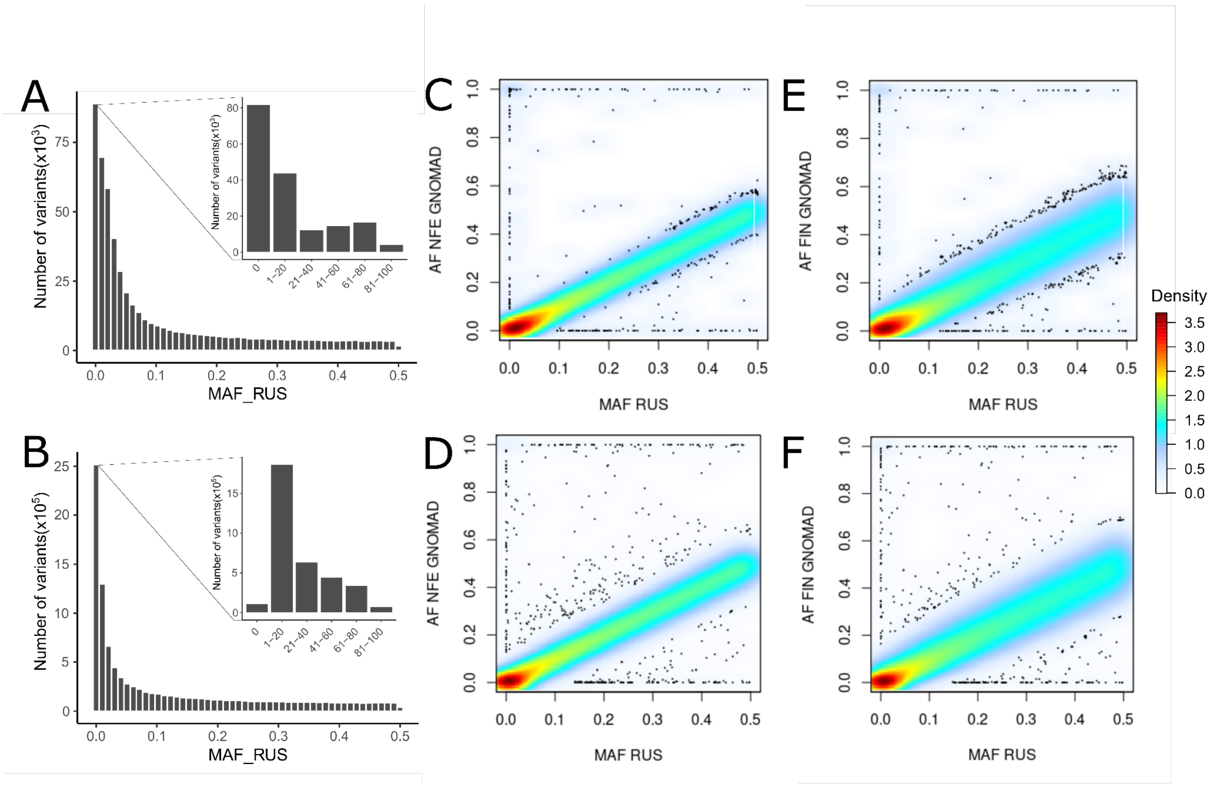


**Supplementary figure S2.** Variant quality control. **(A)** Distribution of MAFs for genotyped variants and allele counts for rare variants (MAF<0.01); **(B)** Distribution of MAFs for all variants and allele counts for rare variants (MAF<0.01); **(C)** comparison of genotyped MAFs in Russian population with allele frequencies in gnomAD Non Finish Europeans; **(D)** comparison of all MAFs in Russian population with allele frequencies in gnomAD Non Finish Europeans; **(E)** comparison of genotyped MAFs in Russian population with allele frequencies in gnomAD Finish samples; **(F)** comparison of all MAFs in Russian population with allele frequencies in gnomAD Finish samples.

### Principal Component Analysis

We used PCA analysis to stratify the Russian population using the PLINK2 tool. Initially we use clumping to find independent variants ($R^2$<0.2). 3,477,939 rare variants with MAF<0.01 were excluded. LD-clumping resulted in 535,727 independent variants ($R^2$<0.2). 4,281 individuals and 535,727 variants were used for PCA (**Sup. Fig. 3A-B**).

We used R library *'adamethods' (v1.2.1)* [35] to iteratively identify PC outliers in PC1-PC4. In each iteration we exclude 3 outliers with the highest distance to 5 nearest neighbors and recalculated clumping variants and PCA. We stopped the procedures at step 45 when max Euclidean distance between two points was minimal **(Sup. Fig. S4)**. Also, we excluded 1 individual from the African population according to further ADMIXTURE analysis. Therefore, the final dataset of PCA analysis included 4,145 individuals and 536,579 clumping variants **(Sup. Fig. 3C-D)**.

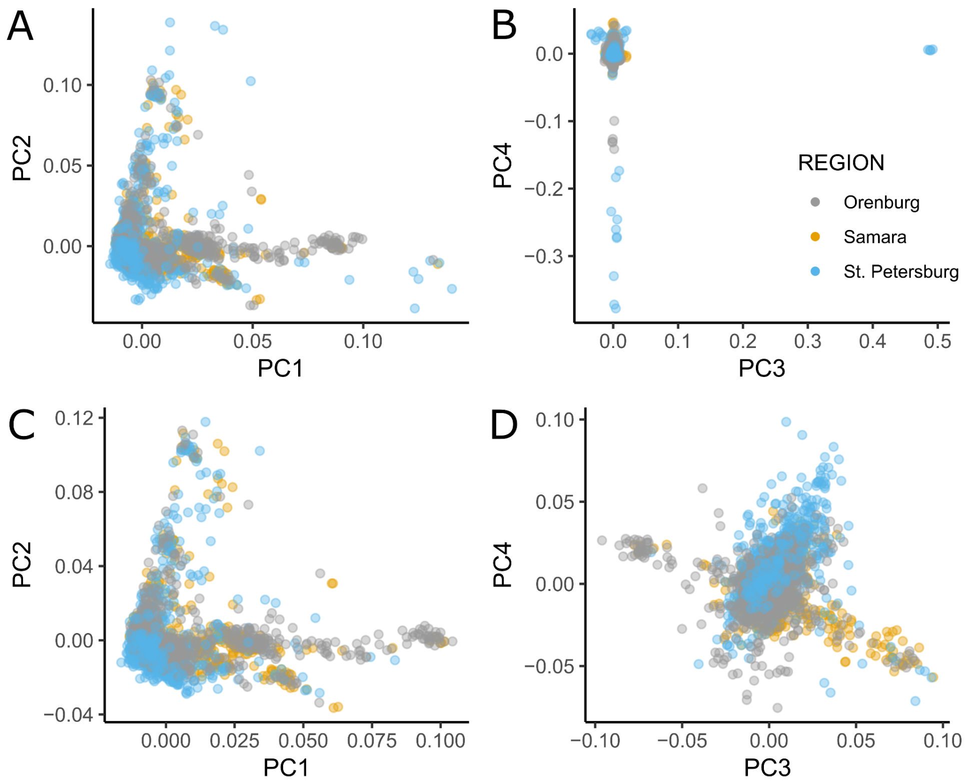


**Supplementary figure S3.** Principal component analysis. **(A)** PC1 and PC2 for Russian population before filtering outliers; **(B)** PC3 and PC4 for Russian population before filtering outliers; **(C)** PC1 and PC2 for Russian population after filtering outliers; **(D)** PC3 and PC4 for Russian population after filtering outliers

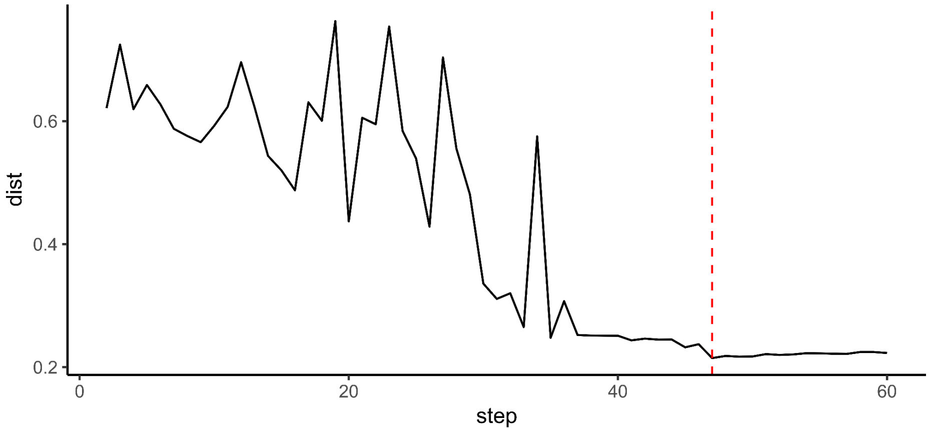


**Supplementary figure S4.** Maximum Euclidean distance in each iteration of filtering outliers

### Comparison of Russian samples with 1000 Genomes

Using python3.6 *'hail'* package we combined the Russian initial dataset (4,281 individuals and 11,077,392 variants) with 1000 Genomes WGS data to contain 10,267,739 variants and 2,504 individuals. We filtered variants with MAF <0.01 and HWE > $1x10^{-4}$ and used clumping ($R^2$<0.2). 506,617 pruned variants and 6,785 individuals were used to build PCs (**Sup. Fig. S5A**). PCA without 136 Russian outliers are shown at **Fig. 2B**. The similar procedure was done with 1000 Genomes Europeans. Combined genotyping data contained 10,267,739 variants and 4,784 individuals. 515,649 pruned variants were used to build PCs. We united central Europeans with British Europeans as they are genetically close. Also, we united Spain and Italy populations (**Sup. Fig. S5B**). PCA without 136 Russian outliers is shown at **Fig. 2C**.

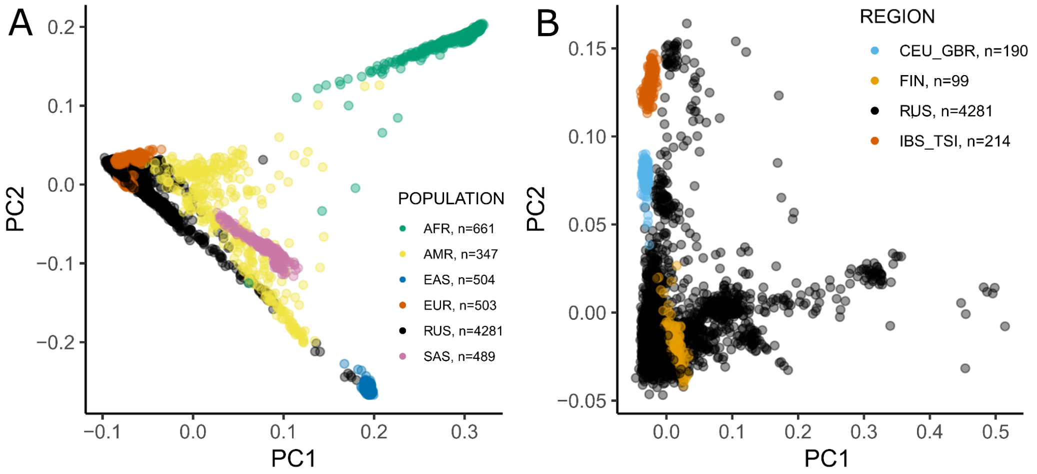


**Supplementary figure S5.** Principal component analysis of Russian samples combined with 1000G without filtering Russian outliers. **(A)** PC1 and PC2 for combined Russian 1000 Genomes data; **(B)** PC1 and PC2 of Russian samples combined only with 1000 Genomes Europeans.

### Clustering of Russian samples

We used the '*SVDFunctions*' *(v1.2)* [36] R package to identify clusters in the Russian population. We selected the number of clusters in such a way that they contained more than 100 individuals and were interpretable in the first 2 principal component space **(Fig. 2D)**. The resulting dendrogram with 6 clusters is shown at **Sup. Fig. S6A** and cluster enrichment among geographical regions are shown at **Sup. Fig. S6B**.

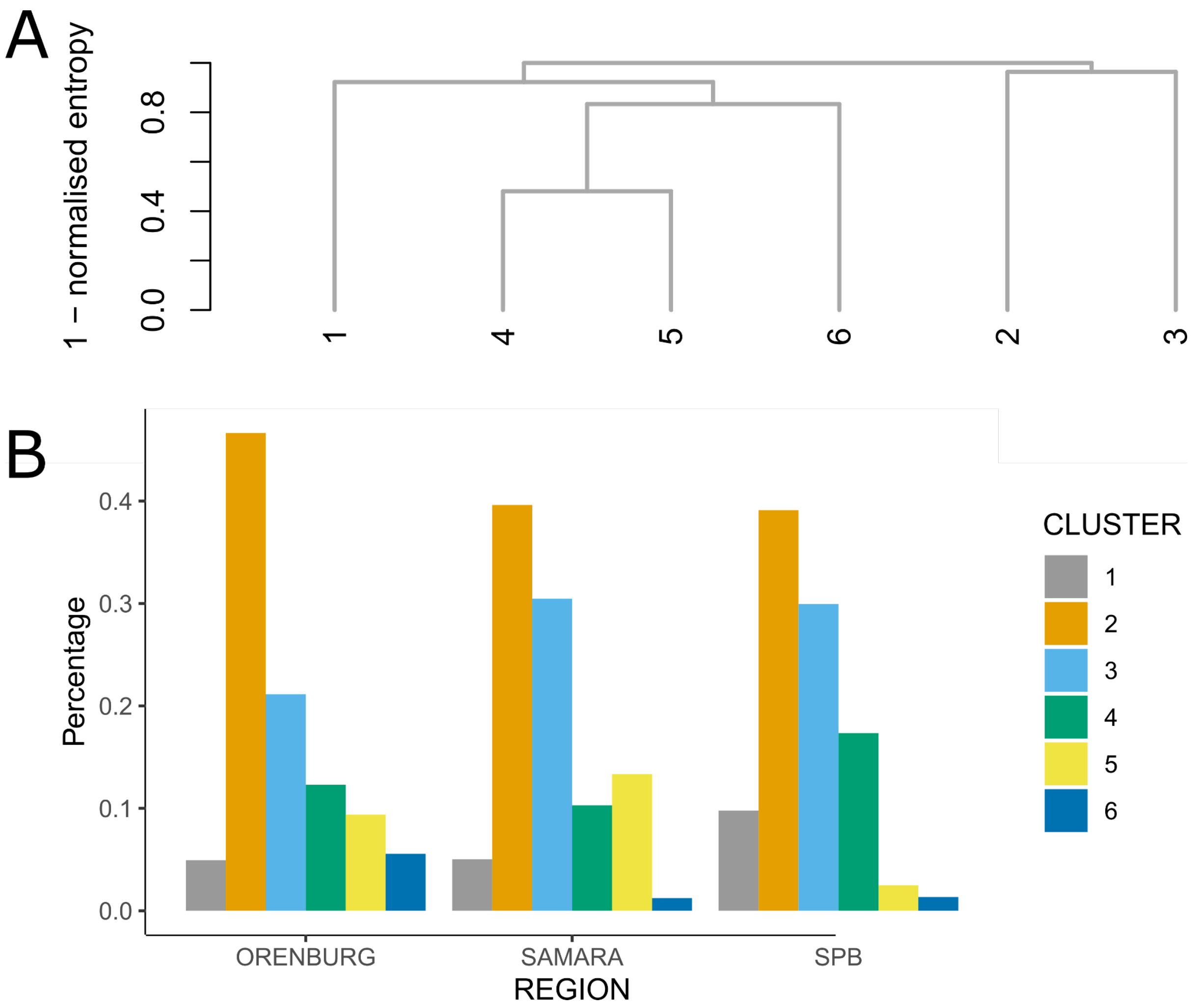


**Supplementary figure S6.** Clustering of Russian samples based on PC1-PC4. **(A)** hierarchy of clusters; **(B)** enrichment of clusters in each geographical region.

### Admixture analysis

To assess the genetic makeup of the Russian population we run ADMIXTURE (v1.3.0) [37] in supervised mode for a LD-pruned combined (ESSE/Starvation Controls -1000 Genomes) dataset with 506,617 pruned variants and 6,649 samples. Initially we identified an additional 14 relatives in the 1000 Genomes dataset using the PLINK2 tool and excluded all 361 related individuals from analysis. Training dataset included 8 populations form 1000 Genomes: African (AFR, N=652), American (AMR, N=347), Vietnam (CDX_KHV, N=192), Central Europeans and British (CEU_GBR, N=190), Chinese and Japanese (CHB_CHS_JPT, N=312), Finish (FIN, N=99), South Europeans (IBS_TSI, N=214) and South Asians (SAS, N=484). The percentage of each training population was determined for each individual in the Russian dataset and arranged by PC1 **(Fig. 2E)**. Aggregated means of each training population for each cluster are shown on **Sup. Fig. S7.**

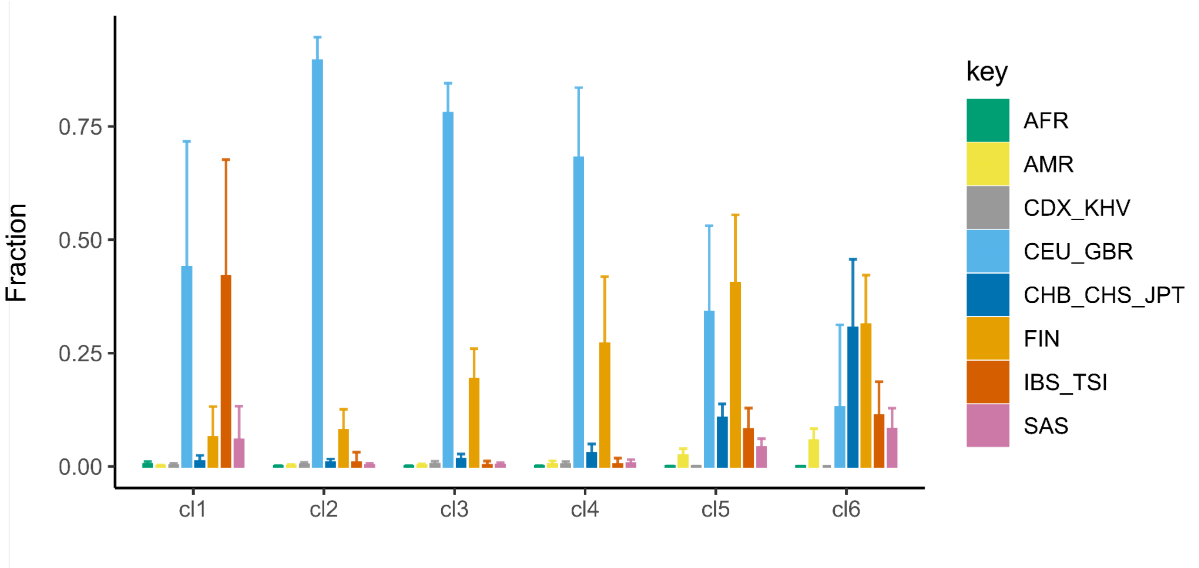


**Supplementary Figure S7.** Mean fractions of each analyzed population by clusters.

### Identity-by-descent (IBD) estimation

We used BEAGLE 4.0 (beagle.r1399.jar) [29] to calculate IBD-sharing statistics between each pair of individuals in the combined (ESSE/Starvation Controls - 1000 Genomes) LD-pruned dataset. We collected only IBD regions with ibdlod quality score more than 3. Then we used ‘merge-ibd-segments.17Jan20.102.jar’ to merge IBD segments if the gap between segments had at most one discordant homozygote and that is less than 0.6 cM in length. We excluded all 361 related individuals from analysis and calculated the total length of all IBD-segments (cM) in each pair of individuals and then median length across each pair of populations. We calculated Euclidean distances to make clasterization and used ‘*gplots’* (v3.0.3) [38] to visualize heatmap matrix (**Fig. 2F).** IBD-searing statistics between each 1000 Genomes population and Russian clusters is shown at **Sup. Fig. S8**

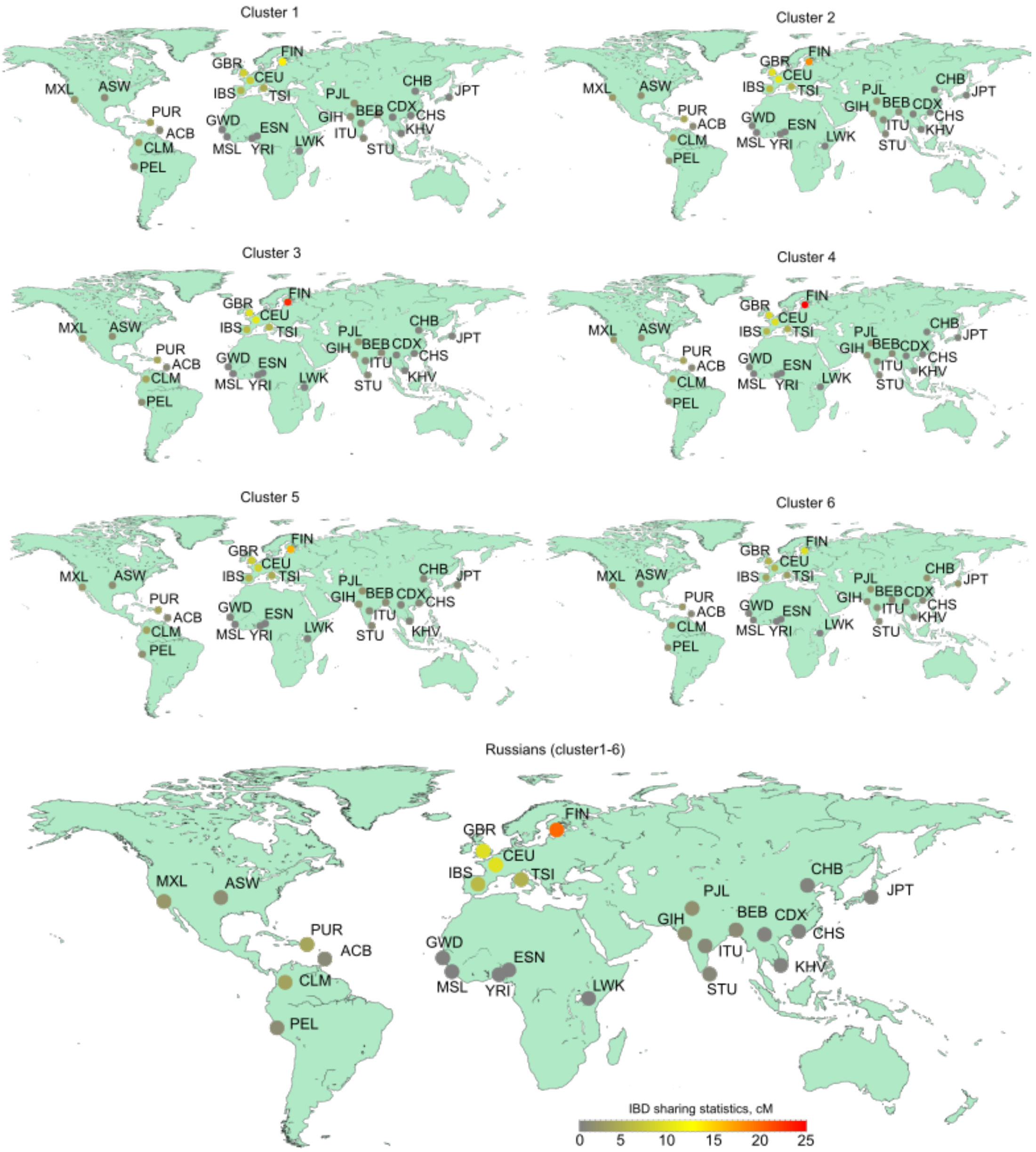


**Supplementary Figure S8.** Mean IBD length between Russians (cl1-cl6) and 1000 genomes populations

### Estimated population size

Estimated population size was calculated for resulting IBD regions with length more than 2cM using IBDne (ibdne23Apr20.ae9.jar) tool [39] for Finnish and Russian populations (**Fig. 3C**). Estimated population size for pairs of clusters in comparison with Finnish populations are shown at **Sup. Fig. S9.**

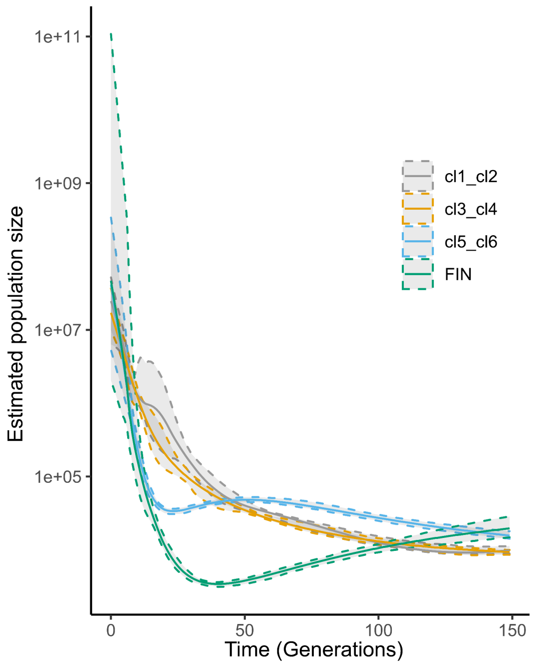


**Supplementary Figure 9.** Estimated population size.

### $F_{st}$ estimation

To estimate the relationship between the Russian population and other populations from 1000G we also calculated the Fst metric. First of all, we used VCFtools (v0.1.15) [40] to calculate $F_{st}$ between Russian geographical regions and 1000G populations using a pruned genotype matrix (506,617 variants and 6,649 individuals). We excluded 361 related individuals and calculated mean $F_{st}$ for each chromosome independently. Distribution of resulting Fst values is shown at **Sup. Fig. S10**. Also, we calculated the Fst between each Russian cluster and each 1000G population. Means across all chromosomes are shown at **Sup. Fig. S11**.

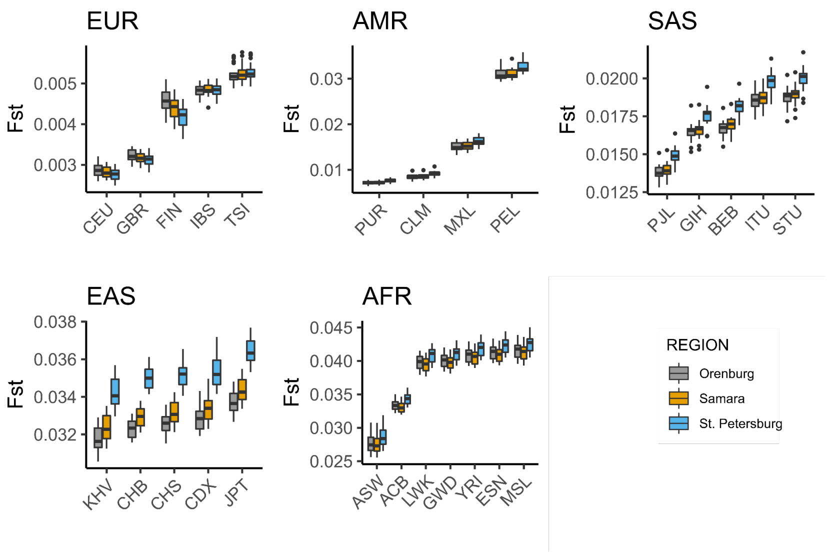


**Supplementary figure S10.** Fst estimation between each region and 1000 Genomes populations

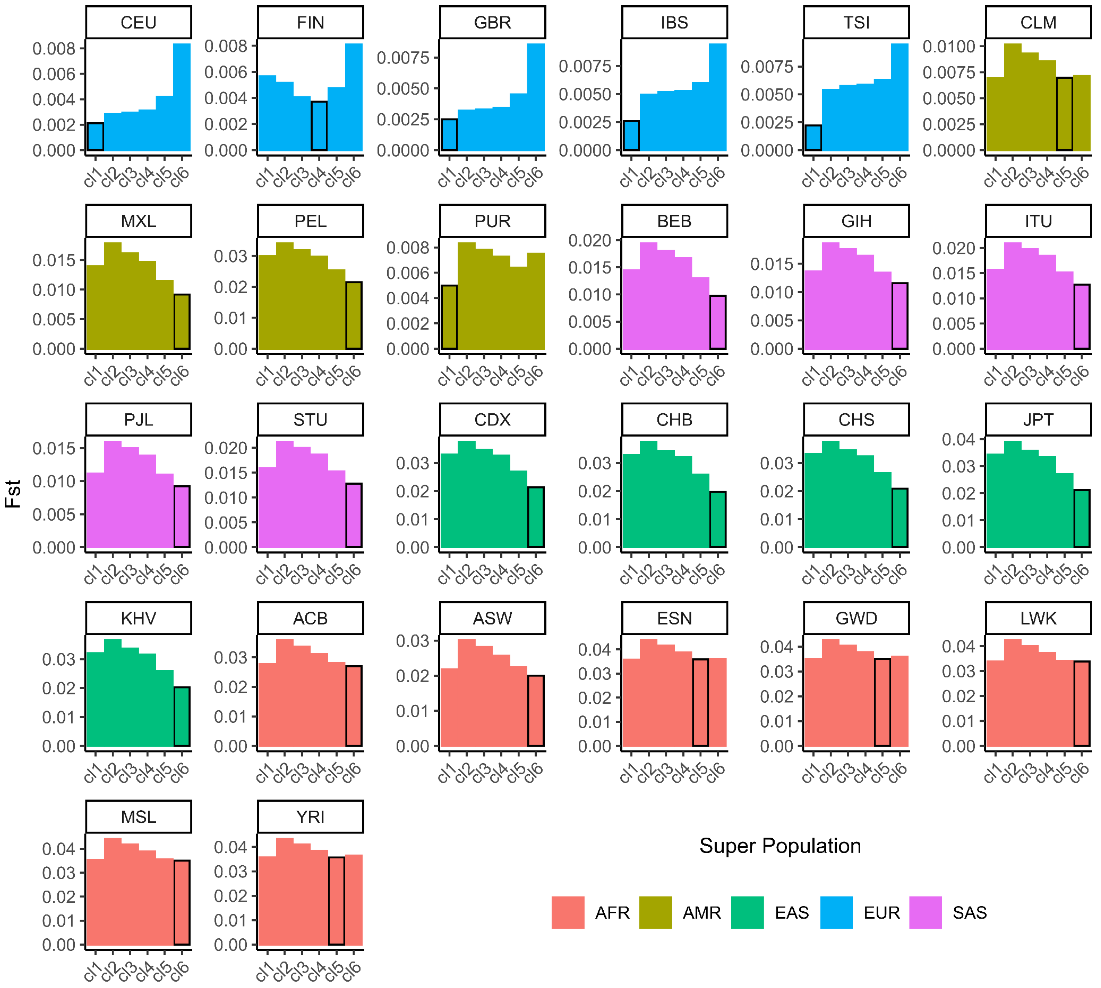


**Supplementary Figure S11.** Fst estimation between each Russian population cluster and 1000 Genomes populations

### Enrichment of Finnish variants

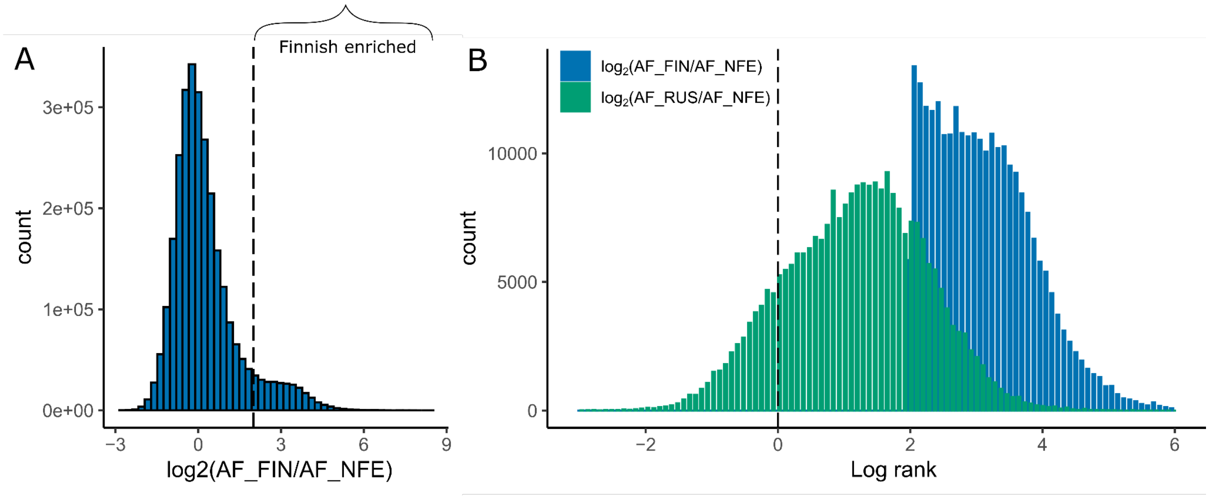


**Supplementary figure S12.** Finnish enriched variants. **(A)** logarithmic ratio of frequencies in Finnish to non-Finnish Europeans for all Russian variants; **(B)** comparison of Finnish enriched variant frequencies in the Russian population.

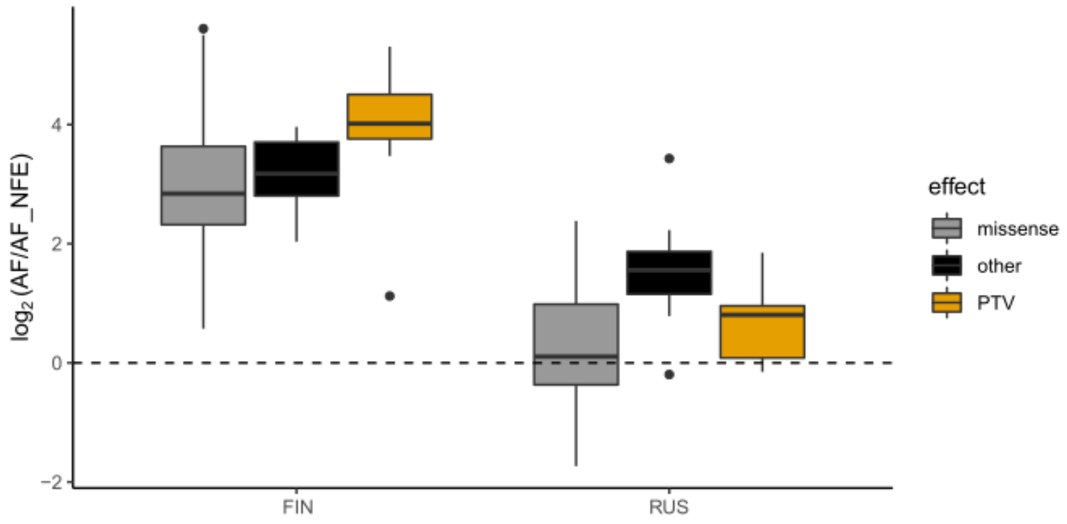


**Supplementary figure S13**. Previously reported Finnish enriched variants that were associated with clinical phenotypes [27].

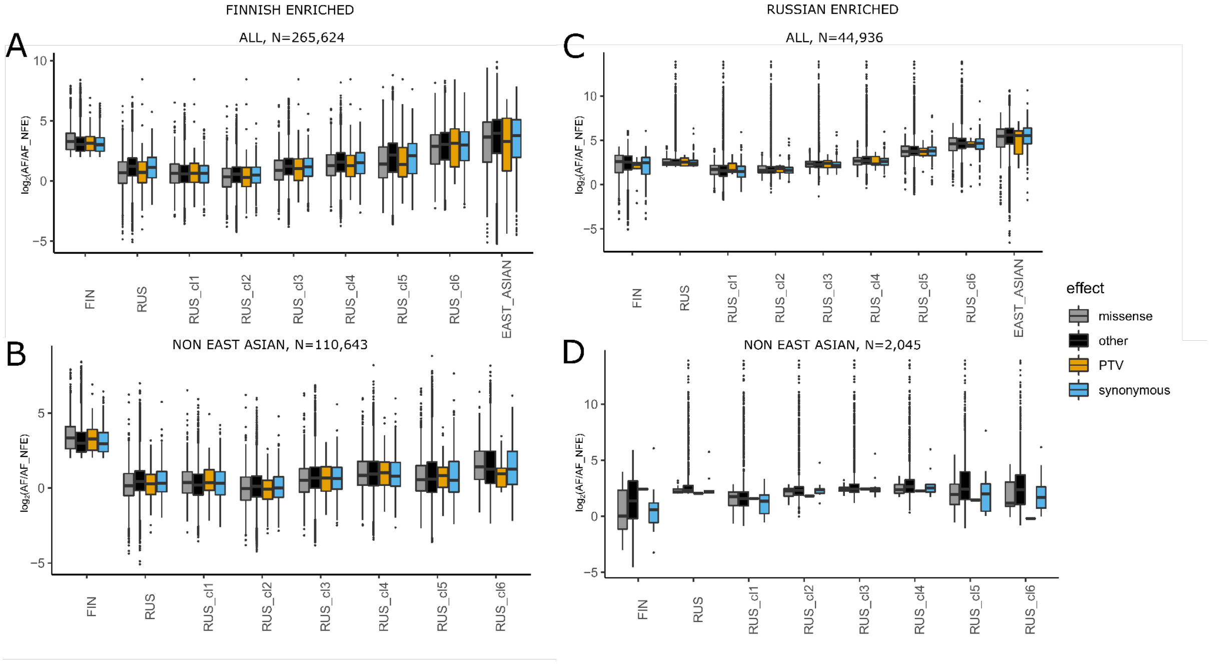


**Supplementary figure S14**. Finnish and Russian enriched variants. **(A)** All Finnish enriched variants; **(B)** Non east Asian Finnish enriched variants; **(C)** All Russian enriched variants; **(D)** Non east Asian Russian enriched variants

### Population Tree Construction

6,288 unrelated Russians and 1000 Genomes individuals were used to construct the maximum likelihood tree based on the combined (ESSE/Starvation Controls - 1000 Genomes) LD-pruned dataset. Additionally we included several populations from the Estonian Genome Diversity Panel (EGDP): **Caucasians** (*Abkhazians, Avars, Azerbaijanis, Balkars, Circassians,*

*Georgians, Kabardins, Kumyks, Lezgins, North-Ossetians, Tabasarans*, N=33), **Europeans** (**East** (*Poles, Belarusians, Ukrainians_east, Ukrainians_north, Ukrainians_west, Cossacks, Cossacks_Kuban, Russians-Central, Russians-North, Russians-West,Russians*): N=25; **North** (*Swedes, Estonians, Finnish, Latvians, Lithuanians, Saami, Ingrians, Karelians, Vepsas, Mordvins*): N=33; **South & West** (*Germans, Hungarians, Moldavians, Albanians, Roma, Croats*): N=17; **Volga Ural** (*Maris, Udmurds, Komis, Tatars, Mishar-Tatars, Kryashen-Tatars, Bashkirs, Chuvashes*): N=25), **Siberians** (**West** (*Mansis, Khantys, Forest-Nenets, Tundra-Nenets, Selkups, Kets*): N=18; **Central** (*Nganasans, Sakha, Evens_Sakha, Evens_Magadan, Evenks, Yakuts*): N=31; **South** (*Shor, Altaians, Tuvinians, Buryats, Mongolians*): N=34, **Northeast** (*Koryaks, Chukchis, Eskimo*), N=25) and **Central Asians** (Rushan-Vanch, Shugnan, Tajiks, Yaghnobi, Turkmens, Uzbek, Kyrgyz_Tdj, Kyrgyz, Kazakhs, Uygurs, Ishkasim): N=24) [41]. Also we added a Russian population (N=25) from the Human Genome Diversity Panel [42]. PCA is shown at **Sup. Fig. S15**. Populational tree based on allele counts for each population was constructed using TreeMix (v.1.12) splitting the genome into blocks with length 500 SNP and using default bootstrapping (**Fig. 3D**) [43].

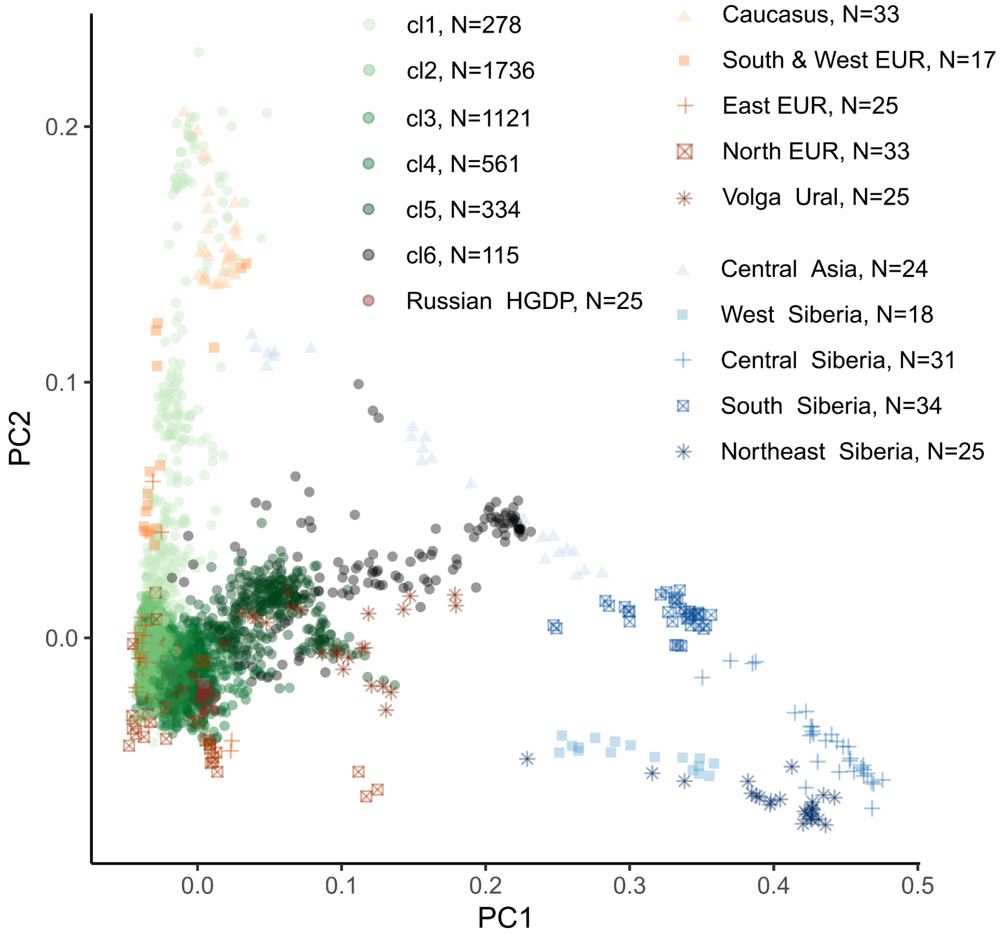


**Supplementary figure S15.** PCA with Russians and several neighboring populations from EGDP.

Finally we run ADMIXTURE analysis in unsupervised mode with 8 genetic clusters and all (N=7,744) unrelated individuals from the Russian dataset, 1000G, HGDP and EGDP. Only local ethnicities show at **Sup. Fig. S16.**

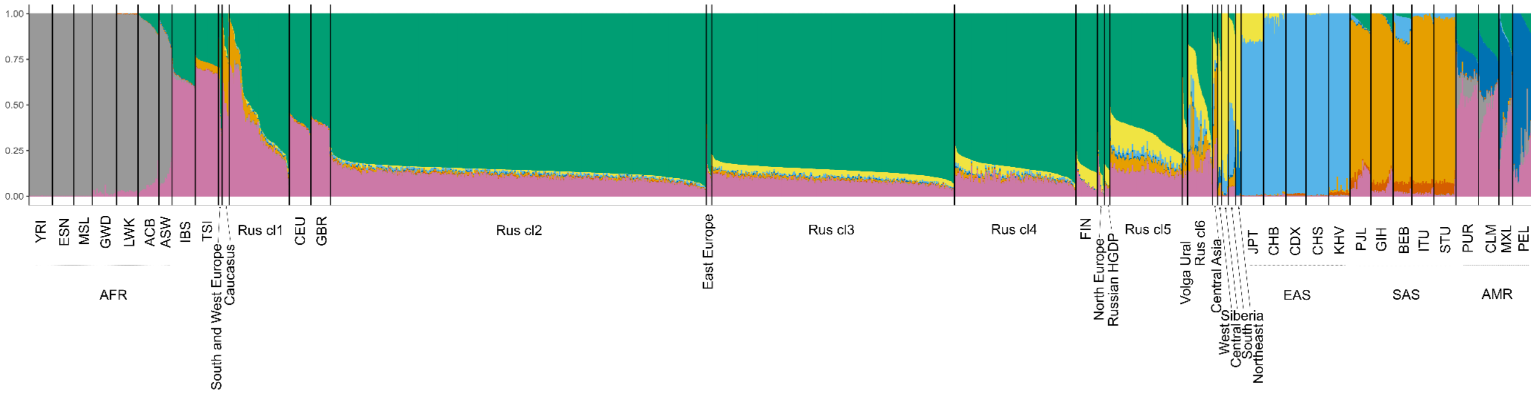


**Supplementary figure S16.** ADMIXTURE analysis in unsupervised mode with 8 clusters.

### GWAS

We performed GWAS for 465 phenotypes. We used linear models for continuous and categorical phenotypes and logistic models for binary phenotypes. Each GWAS model was adjusted for age, sex and PC1-PC4 and created using Python3.6 *'hail'* package. Since some phenotypes were not determined for all individuals, additional quality control (MAF>0.01, HWE>0.0001) was performed separately for each GWAS. First, we checked well-known associations for cholesterol (N=3,856) and blood uric acid levels (N=3,457) to ensure the quality of the phenotyping. We found a strong association of the rs7412 variant (gene *APOE,* beta = -0.43, p=4.7x$10^{-29)}$ and rs4970834 (gene *CELSR2,* beta= -0.15 ,p=1.81x$10^{-8}$) with LDL levels. rs7412 and rs4970834 are known variants associated with LDL levels in UK Biobank (beta = -0.51, p=0) and (beta=-0.14,p=0), correspondingly (**Sup. Fig. 17A**). For blood uric acid we found leading variants rs4697701 (gene *SLC2A9,* beta = -22.2, p=2.21x$10^{-28)}$ and rs45499402 (gene *ABCG2,* beta = 21.9, p=3.99x$10^{-12}$) that are replicated in UK Biobank (beta = -22.7, p=0) and (beta=14.1,p=0), correspondingly (**Sup. Fig. 17B**).

Moreover, we confirmed an association of rs13266066 (AF RUS=0.4358, AF NFE = 0.4763, AF FIN = 0.4397) with smoking initiation that was previously predicted by MTAG algorithm (beta = -0.007, p=1 x $10^{-10}$, beta was reversed to match models), while initial association was not strong (beta = -0.012, p=0.0037, beta was reversed to match models) [44]. In our data we found that the allele frequency of rs13266066 is significantly lower in a group of individuals who never smoked (N never smoked = 2,391; AF never smoked = 0.414, N controls = 1,488; AF controls = 0.475, beta = -0.28, p=3.74x$10^{-8}$) (**Sup. Fig. 17C)**. In the UK biobank rs13266066 is nominally negatively associated with past tobacco smoking, never smoking and age of smoking initiation (p=1.5x$10^{-7}$, p=2x$10^{-4}$,p=3x$10^{-4}$), correspondingly.

To reduce the possibility of technical artifacts associated with this observation we looked at only directly genotyped variants in this locus and confirmed the presence of highly-associated rs11781072 (p=1.45x$10^{-6}$). eQTL properties of rs13266066 increased expression of *PTK2* gene in a cerebrum (p=8.8x$10^{-12}$).

We made gene prioritization with GPrior [45] for *PTK2* using 251 genes associated with trait 'Smoking initiation (ever regular vs never regular)' (p<6x$10^{-6}$) from GWAS catalog as training set and GWAS summary statistic 'Smoking never' from UK Biobank (phenocode:

20116_0). We used only variants with p value less than $1x10^{-3}$ that were initially annotated by genes with POSTGAP and then prioritized with GPrior (**Sup. Fig. S18**). The results of prioritization showed that the PTK2 was most likely associated with never smoking status according to all prediction models except SVM.

We also found an association rs7972723 (AF RUS=0.1465, AF NFE = 0.1447, AF FIN = 0.1680) with the current smoking status (N current smoker = 834, AF current smoker = 0.189, N controls = 3,045; AF controls = 0,137, beta = 0.43, $p=2.08x10^{-8}$) (**Sup. Fig. 17D**). Current smokers included both people who are currently smoking and who quit smoking less than 1 year ago. Interestingly, allele frequency of rs7972723 increased in three groups of smoking status: never smoked - 0.136, smoking in the past - 0.141, current smoker - 0.189 ($p=1.63x10^{-7}$). eQTL properties of rs7972723 increase expression of *ACSM4* gene in adipose subcutaneous tissue ($p=1.7x10^{-7}$). rs7972723 is nominally replicated in UK Biobank as associated with smoking in the past (beta=0.0036, p=0.02). Significantly associated directly genotyped variant rs7953422 also presented in this locus ($p=3.54x10^{-8}$). We didn't perform gene prioritization for ACSM4 due to a small training gene set for smoking cessation (N=21).

In addition, we found some new associations. For example variant rs56046524 (AF RUS=0.3537, AF NFE = 0.3938, AF FIN = 0.3632) associated with abdominal obesity (N cases=1,405; AF cases = 0.306, N controls = 2,462; AF controls=0.378, beta=-0.324, $p=3.7x10^{-9}$) (**Sup. Fig. 17E**). Also variant rs11948871 (AF RUS=0.2024, AF NFE =0.1727, AF FIN = 0.1479) is associated with increased blood pressure in the second half of pregnancy (N cases=366; AF cases=0.279, N controls = 1,642; AF controls = 0.185, beta=0.55,$p=1.4x10^{-8}$) (**Sup. Fig. 17F**). Both these observations are accompanied with highly associated directly genotyped variants (rs100016, $p=4.69x10^{-7}$; rs6872733, $p=2.21x10^{-5}$), correspondingly.

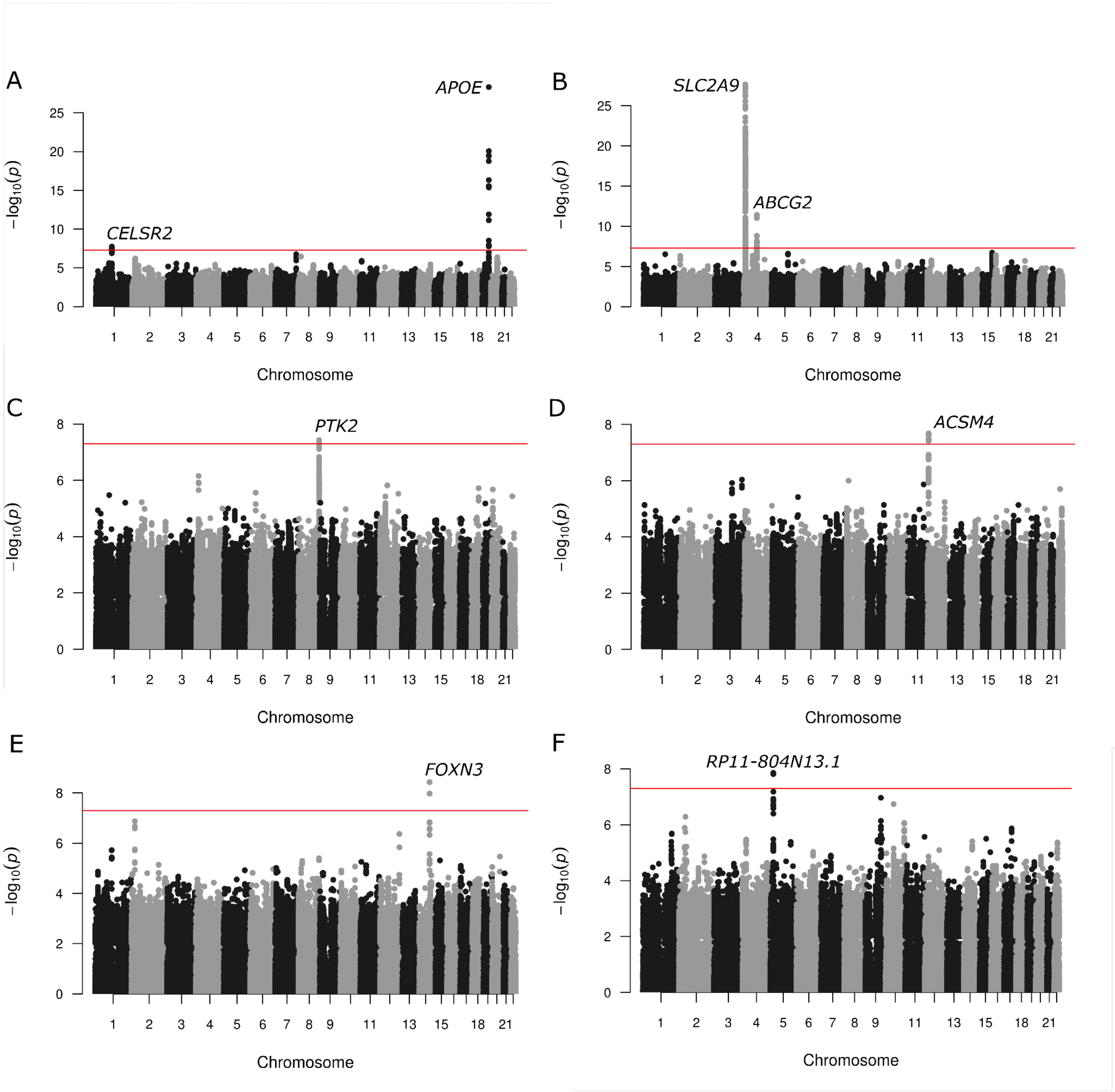


**Supplementary figure S17.** GWAS. **(A)** LDL-levels; **(B)** Uric acid levels; **(C)** Smoking (never); **(D)** Smoking (current); **(E)** Abdominal obesity; **(F)** Increased blood pressure in pregnant women

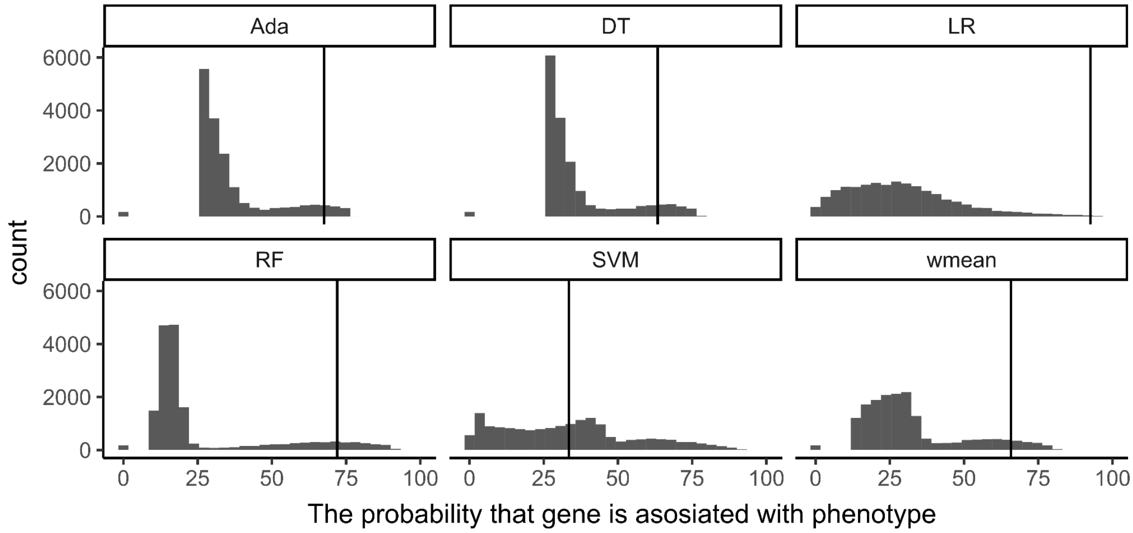


**Supplementary figure S18.** The results of PTK2 gene prioritization with Never smoke phenotype. The black line is the value for PKT2 gene

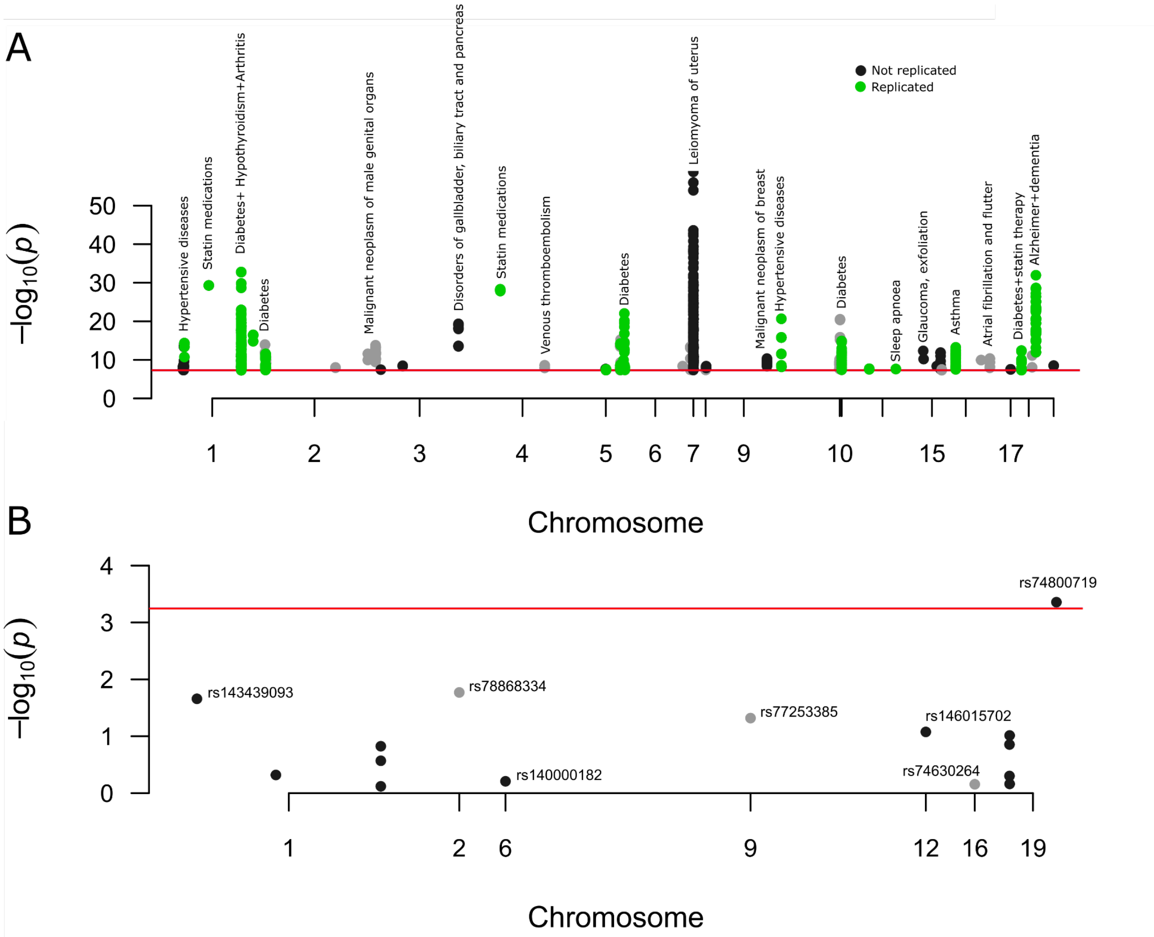


**Supplementary figure S19.** Replication of genome-wide significant Finnish enriched variants (MAF RUS > 0.01, log-ratio EAS NFE < 2) in Russian biobank; **(A)** All genome-wide significant Finnish enriched variants (MAF RUS > 0.01, log-ratio EAS NFE < 2); **(B)** Replication of clumped variants united in 8 phenotypes
